# Potent neutralization of SARS-CoV-2 variants of concern by an antibody with a unique genetic signature and structural mode of spike recognition

**DOI:** 10.1101/2021.05.16.444004

**Authors:** Kevin J. Kramer, Nicole V. Johnson, Andrea R. Shiakolas, Naveenchandra Suryadevara, Sivakumar Periasamy, Nagarajan Raju, Jazmean K. Williams, Daniel Wrapp, Seth J. Zost, Clinton M. Holt, Ching-Lin Hsieh, Rachel E. Sutton, Ariana Paulo, Edgar Davidson, Benjamin J. Doranz, James E. Crowe, Alexander Bukreyev, Robert H. Carnahan, Jason S. McLellan, Ivelin S. Georgiev

## Abstract

The emergence of novel SARS-CoV-2 lineages that are more transmissible and resistant to currently approved antibody therapies poses a considerable challenge to the clinical treatment of COVID-19. Therefore, the need for ongoing discovery efforts to identify broadly reactive monoclonal antibodies to SARS-CoV-2 is of utmost importance. Here, we report a panel of SARS-CoV-2 antibodies isolated using the LIBRA-seq technology from an individual who recovered from COVID-19. Of these antibodies, 54042-4 showed potent neutralization against authentic SARS-CoV-2 viruses, including variants of concern (VOCs). A cryo-EM structure of 54042-4 in complex with the SARS-CoV-2 spike revealed an epitope composed of residues that are highly conserved in currently circulating SARS-CoV-2 lineages. Further, 54042-4 possesses unique genetic and structural characteristics that distinguish it from other potently neutralizing SARS-CoV-2 antibodies. Together, these findings motivate 54042-4 as a lead candidate for clinical development to counteract current and future SARS-CoV-2 VOCs.

## Introduction

The COVID-19 pandemic caused by a novel coronavirus from the *Sarbecovirus* genus, SARS-CoV-2, spawned an unprecedented global research effort dedicated to therapeutic countermeasure development resulting in rapid US FDA Emergency Use Authorization (EUA) for vaccines and monoclonal antibodies (Jones et al., 2021; Weinreich et al., 2021). The primary target for vaccine and antibody therapeutic development is the SARS-CoV-2 spike (S) protein, which facilitates host-cell attachment and entry (Wrapp et al., 2020). The emergence of distinct viral lineages that accumulate substitutions in the S protein pose a significant threat to the countermeasures currently approved for clinical use (Chen et al., 2021; Wang et al., 2021). Continued genomic surveillance and persistent efforts to identify novel antibodies with distinct binding modes and mechanisms of action are crucial to maintain availability of therapeutics in the event of further neutralization-escape by SARS-CoV-2 variants of concerns (VOCs).

SARS-CoV-2 spike is a class I viral fusion protein that is a trimer of heterodimers composed of S1 and S2 subunits (Wrapp et al., 2020). S1 initiates attachment to the receptor angiotensin-converting enzyme 2 (ACE2), whereas S2 drives membrane fusion by refolding from a prefusion to postfusion conformation (Li, 2016; Tortorici and Veesler, 2019). The primary contact of ACE2 and spike is in the receptor-binding domain (RBD) of the S1 subunit, which is composed of a receptor binding motif (RBM) and RBD core. The three RBDs within each spike can adopt an ACE2-accessible “up” conformation and an ACE2-inaccessible “down” conformation via a hinge-like motion (Shang et al., 2020). As a result, although multiple neutralizing epitopes on spike have been identified (Brouwer et al., 2020; Chi et al., 2020; Suryadevara et al., 2021; Zost et al., 2020), the RBD serves as the dominant target of neutralizing antibodies via antagonism of ACE2 binding (Piccoli et al., 2020).

Neutralizing antibodies targeting the RBD have been characterized extensively and partition into different classes based on binding mode, ACE2 interface overlap, and cross-reactivity with other *Sarbecoviruses*. For example, neutralizing antibodies predominantly encoded by IGHV3-53 and IGHV3-66 have epitopes directly covering the ACE2 interaction footprint in the RBM (Yuan et al., 2020a). Examples of this class of antibodies are clinical EUA candidates REGN10933 and COV2-2196 (Hansen et al., 2020; Zost et al., 2020). Antibodies that bind the RBM but are more distal to the ACE2 interface form another distinct class that includes REGN10987 and COV2-2130 (Dong et al., 2021; Hansen et al., 2020). Additionally, antibodies such as CR3022 and ADG-2 that cross-react with other coronaviruses comprise a more diverse group that target conserved residues in the RBD-core (Pinto et al., 2020; Wec et al., 2020; Yuan et al., 2020b).

The continued transmission of SARS-CoV-2 in the human population has led to the evolution of VOCs with increased transmissibility and resistance to available medical countermeasures (Alpert et al., 2021; Kuzmina et al., 2021). Some of the most consequential amino acid substitutions observed so far have occurred in the RBD, particularly N501Y in the B.1.1.7, B.1.351, and P.1 lineages, and the additional combination of K417N/T and E484K in the P.1 and B.1.351 lineages. In particular, N501Y is thought to increase affinity for ACE2 (Starr et al., 2020) potentially resulting in increased infectivity, whereas E484K disrupts the antigenic landscape of the RBD that can lead to substantial decreases in neutralization titers (Hoffmann et al., 2021; Wang et al., 2021). In some cases, SARS-CoV-2 VOCs also escape neutralization by polyclonal antibodies in the serum from vaccine recipients and individuals previously infected with SARS-CoV-2 (Chen et al., 2021; Wang et al., 2021). These observations highlight the critical need for a wide range of potently neutralizing antibodies insensitive to substitutions arising in VOCs.

To address this challenge, we applied LIBRA-seq, a recently developed antibody-discovery technology (Setliff et al., 2019; Shiakolas et al., 2020), to interrogate the B cell repertoire of an individual who had recovered from COVID-19. Our efforts led to the discovery of a potently neutralizing antibody, designated 54042-4, which uses a unique genetic signature and structural mode of SARS-CoV-2 RBD recognition to maintain neutralization potency to known VOCs. Antibody 54042-4 therefore may serve as a viable candidate for further prophylactic or therapeutic development for protection against a broad range of SARS-CoV-2 variants.

## RESULTS

### Identification of SARS-CoV-2-neutralizing antibodies by LIBRA-seq

To identify SARS-CoV-2 S-directed antibodies, we utilized LIBRA-seq (Linking B Cell receptor to antigen specificity through sequencing), a technology that enables high-throughput simultaneous determination of B cell receptor sequence and antigen reactivity at the single-cell level, expediting the process of lead candidate selection and characterization (Setliff et al., 2019). The LIBRA-seq antigen-screening library included SARS-CoV-2 spike stabilized in a prefusion conformation (Hsieh et al., 2020), along with antigens from other coronaviruses and negative-control antigens. Antigen-specific B cells were isolated from a donor with potently neutralizing antibodies in serum (1:258 NT_50_) three months after infection confirmed by nasal swab RT-PCR testing for SARS-CoV-2 (**Supplemental Figure 1A**). Of the 73 IgG^+^ B cells with high LIBRA-seq scores (≥1) for SARS-CoV-2 S (**Figure 1A, Supplemental Figure 1B**), we chose nine lead candidates with diverse sequence characteristics, CDRH3 length, and germline V gene usage for characterization as recombinant monoclonal antibodies (**Figure 1B**). Binding to SARS-CoV-2 S by ELISA was confirmed for eight of these antibodies, with the only exception being antibody 54042-2, in agreement with its lower LIBRA-seq score (**Figure 1B, Supplemental Figure 1C**). Five of these antibodies showed SARS-CoV-2 neutralization activity in a high-throughput neutralization screen using a live chimeric VSV displaying SARS-CoV-2 spike protein (Case et al., 2020) (**Figure 1B**). Full dose-response neutralization curves in the chimeric VSV assay were obtained for four of these five antibodies, with antibody 54042-4 showing the best potency, at a half-maximal inhibitory concentration (IC_50_) of 9 ng/mL (**Figure 1C**).

**Figure 1.**
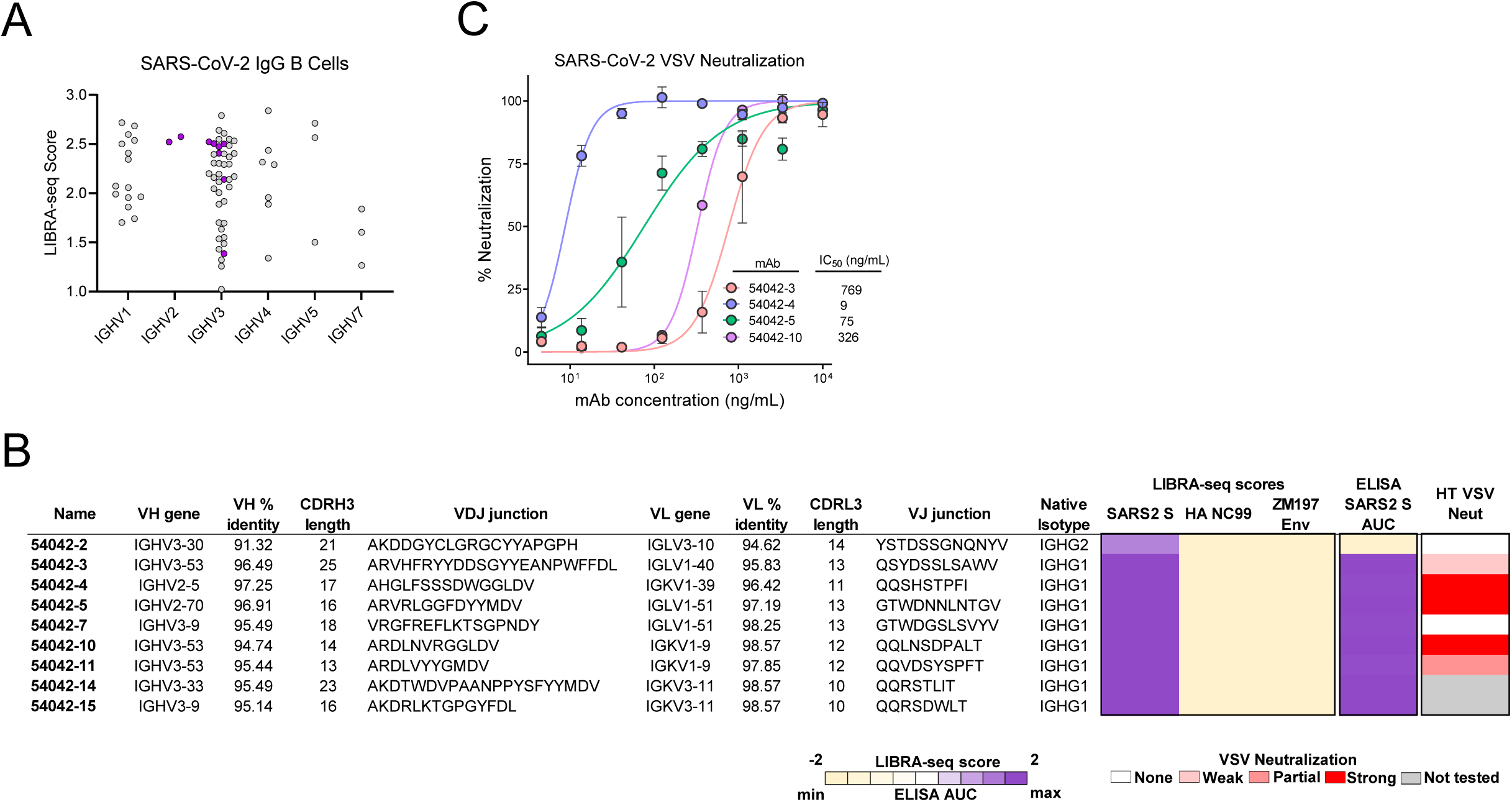
Identification and characterization of SARS-CoV-2 antibodies isolated using LIBRA-seq. **(A)** Variable heavy gene usage (x-axis) as a function of IgG^+^ B cells with a SARS-CoV-2 spike LIBRA-seq score (>1) (y-axis). The nine lead candidates are highlighted in purple. **(B)** RTCA VSV-SARS-CoV-2 neutralization by lead candidate antibodies. IC_50_ values are calculated by non-linear regression analysis by GraphPad Prism software. **(C)** Sequence characteristics and antigen specificity of nine lead candidate antibodies from a recovered COVID-19 donor. Percent identity is calculated at the nucleotide level, and CDR length and sequences are displayed at the amino acid level. LIBRA-seq scores for each antigen are displayed as a heatmap with a LIBRA-seq score of -2 displayed as light yellow, 0 as white, and 2 in purple; in this heatmap scores lower or higher than that range are shown as -2 and 2, respectively. ELISA binding data for SARS-CoV-2 S are displayed as a heatmap of the AUC analysis calculated from (**Supplemental Figure 1C**) with AUC of 0 displayed as light yellow, 50% maximum as white, and maximum AUC as purple.

### Antibody 54042-4 targets the SARS-CoV-2 receptor-binding domain

Because of the potent (≤ 10 ng/mL) virus neutralization observed for 54042-4, we selected this antibody for further characterization. ELISAs performed with purified RBD, NTD, S1, and S2 proteins revealed 54042-4 IgG bound to the SARS-CoV-2 S1 subunit as well as the RBD (**Figure 2A, Supplemental Figure 2)**. To determine the affinity of the antibody-antigen binding interaction, biolayer interferometry experiments were performed by measuring the association and dissociation kinetics of immobilized 54042-4 IgG binding to a soluble protein comprising the RBD and subdomain-1 (SD1) of the SARS-CoV-2 S protein. Curve-fitting resulted in a calculated *K*_D_ of 21.8 nM (**Figure 2B**). Given the neutralization potency of 9 ng/mL (60 pM), these data suggest that the IgG avidly binds to the S protein on the surface of the virus. To assess whether 54042-4 neutralizes viral infection by directly competing with ACE2, a receptor-blocking assay was performed by testing the competition of 54042-4 with soluble ACE2 for binding to SARS-CoV-2 S. The results demonstrated that 54042-4 inhibits interaction of ACE2 to SARS-CoV-2 S protein, unlike the control antibodies CR3022, an extensively characterized SARS-CoV antibody that binds a cryptic epitope in the RBD (Yuan et al., 2020b), and the influenza hemagglutinin-specific 3602-1707 (Setliff et al., 2019) (**Figure 2C**). Next, we performed competition ELISA to determine if 54042-4 competes for binding with three other RBD-directed antibodies with distinct epitopes. These antibodies included COV2-2196 and COV2-2130, which form the basis of AZD7442, an antibody cocktail currently under investigation in clinical trials for COVID-19 treatment and prevention (ClinicalTrials.gov Identifiers: NCT04625725, NCT04723394, NCT04518410, and NCT04501978) and CR3022. The competition experiment showed that 54042-4 competed for binding to SARS-CoV-2 S protein with COV2-2130, but not COV2-2196 or CR3022 (**Figure 2D**). Together, these results suggest that 54042-4 targets an epitope on SARS-CoV-2 RBD that at least partially overlaps with the binding sites for both ACE2 and other potently neutralizing RBD antibodies.

**Figure 2:**
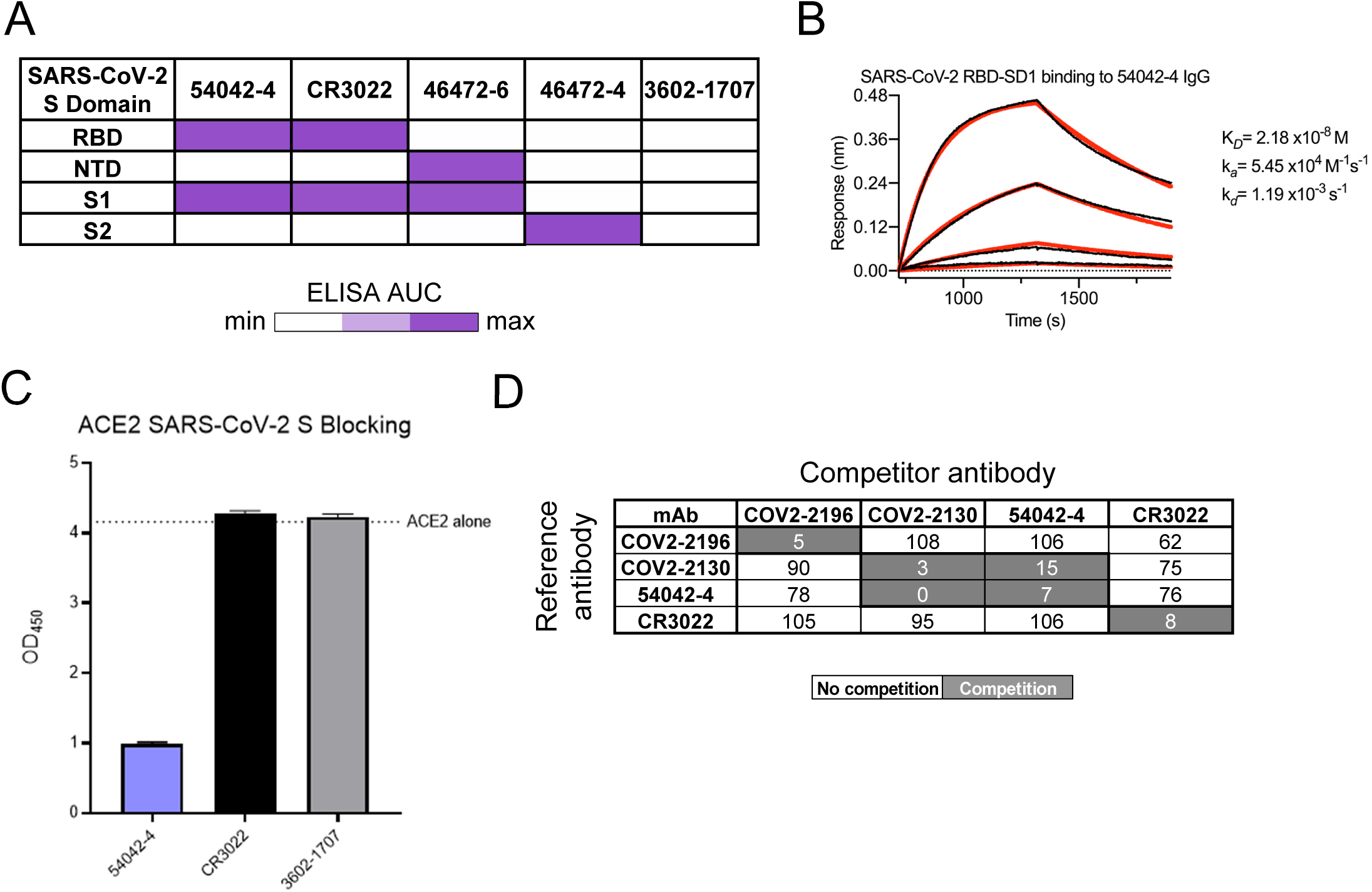
54042-4 functional characterization. **(A)** ELISA binding values against SARS-CoV-2 subdomains are displayed as a heatmap of AUC values calculated from the data in (**Supplemental Figure 2)**. Antibodies CR3022, 46472-6, 46472-4 were used as positive controls for the RBD, NTD, and S2 antigens, respectively (Shiakolas et al., 2020; Yuan et al., 2020b). 3602-1707 was included as an influenza HA-specific negative control antibody (Setliff et al., 2019). **(B)** A biolayer interferometry sensogram that shows binding to recombinant SARS-CoV-2 RBD-SD1. Binding data are depicted by the black lines and the best fit line of the data to a 1:1 binding model is shown in red. **(C)** SARS-CoV-2 spike/ACE2 inhibition ELISA is shown, for 54042-4, SARS-CoV-2 antibody CR3022 and negative control HA-specific antibody 3602-1707. For each antibody, the ACE2 binding signal is depicted on the y-axis, in comparison to ACE2-only binding to SARS-COV-2 spike shown as a dotted line. **(D)** Competition ELISA of 54042-4 with antibodies COV2-2196, COV2-2130, and CR3022. Values in white indicate no competition (presence of competing antibody reduced reference antibody binding by less than 30%) and values in dark grey indicate competition (presence of competing antibody reduced reference antibody binding by more than 60%).

### 54042-4 binds the apex of the SARS-CoV-2 RBD in the down conformation

To gain a better understanding of the recognition of SARS-CoV-2 S by antibody 54042-4, we determined a 2.7 Å resolution cryo-EM structure of the 54042-4 antigen-binding fragments (Fabs) bound to the SARS-CoV-2 S extracellular domain (ECD) modified so that all three RBDs were disulfide-locked in the down conformation (Henderson et al., 2020) (**Figure 3A**). Local refinement of one RBD bound to a 54042-4 Fab was performed to improve the interpretability of the map at the binding interface, resulting in a local 3D reconstruction with a resolution of 2.8 Å (**Figure 3B**). The structure revealed that 54042-4 forms an extensive interface with the RBD, making contacts through the CDRL1, CDRL3, and all three CDRs of the heavy chain, to form a clamp around the apex of the RBM saddle (**Figure 3C,D, Supplemental Figure 3**). The primary interactions involve RBD residues 439–450, with a network of hydrogen bonds between the 54042-4 heavy chain and RBD residues 443–447 (**Figure 3C**). From CDRH3, Ser99 forms a hydrogen bond with RBD residue Ser443, and a hydrogen bond is formed between the mainchain atoms of Phe97 and Val445. From CDRH2, Asp56 forms a hydrogen bond and salt bridge with Lys444, whereas Arg58 forms hydrogen bonds with mainchain atoms from Gly446 and Gly447. The CDRH1 contributes a lone residue, Ile32, to the binding interface, forming minor contacts near Leu441. The 54042-4 light chain surrounds the opposite side of this RBD region, mediating interactions primarily through hydrophobic contacts formed by CDRL1 and CDRL3 near RBD residue Val445 (**Figure 3D**). Additional light chain contacts are made with residues 498–500 of the RBD, including a hydrogen bond between His92 of CDRL3 and Thr500, and hydrophobic interactions involving CDRL1 Phe30 and Tyr32. Notably, the complex structure indicated that a number of spike substitutions associated with current VOCs are unlikely to affect recognition by antibody 54042-4. For example, RBD residue Asn501 (present as Tyr501 in several VOCs, including B.1.1.7, B.1.351, and P.1) lies just outside of the 54042-4 epitope, whereas the Cα atoms of Glu484 (present as Lys484 or Gln484 in, e.g., B.1.351, P.1, and B.1.617) and Leu452 (present as Arg452 in B.1.427) are approximately 18 and 14 Å away from the Cα atoms of the nearest 54042-4 residue, respectively (**Figure 3B**).

**Figure 3:**
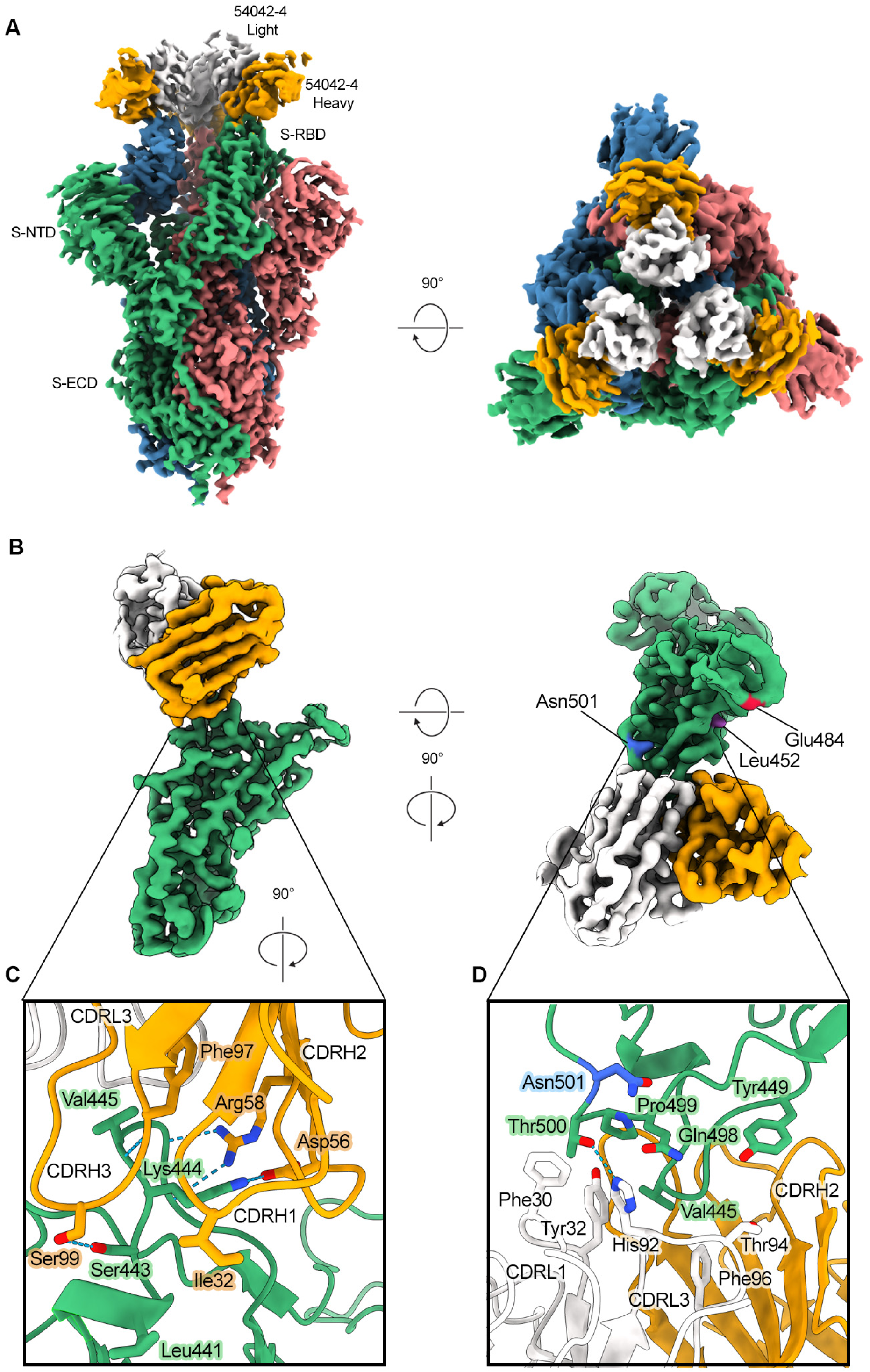
Atomic resolution of 54042-4 binding mode to SARS-CoV-2 S. **(A)** 3D reconstructions of side and top views of Fab 54042-4 bound to SARS-CoV-2 spike. **(B)** Focused refinement maps showing the 54042-4 epitope at the apex of the RBM in the down position (left). Top-down view of the 54042-4 epitope showing heavy and light chain contacts, as well as residues outside of the binding interface that are mutated in circulating VOCs (right). **(C)** The 54042-4 heavy chain binds to RBD residues 443–447 primarily through a network of hydrogen bonds involving CDRH2 and CDRH3 and hydrophobic contacts involving Ile32 of CRDH1. **(D)** The 54042-4 light chain contacts RBD residues 498–500 through a hydrogen bond between Thr500 and His92 of CDRL3 and hydrophobic contacts involving Phe30 and Tyr32 of CDRL1.

### Antibody 54042-4 has a unique genetic signature and structural mode of RBD recognition

Public clonotype sequence signatures (those shared by multiple individuals recovered from COVID-19 infection) have been identified for potently neutralizing SARS-CoV-2 antibodies, including antibodies currently in clinical trials or approved for emergency use (Nielsen et al., 2020; Yuan et al., 2020a). To investigate whether antibody sequences that are closely related to 54042-4 can be identified among known SARS-CoV-2 antibodies, we searched the CoV-AbDab database that contains paired heavy-light chain sequences of coronavirus antibodies (Raybould et al., 2021). Notably, antibodies with high sequence identity to the 54042-4 CDRH3 and CDRL3 were not identified, whether or not the search was restricted to the *IGHV2-5* heavy chain and *IGKV1-39* light chain genes utilized by 54042-4 (**Figure 4A**).

**Figure 4:**
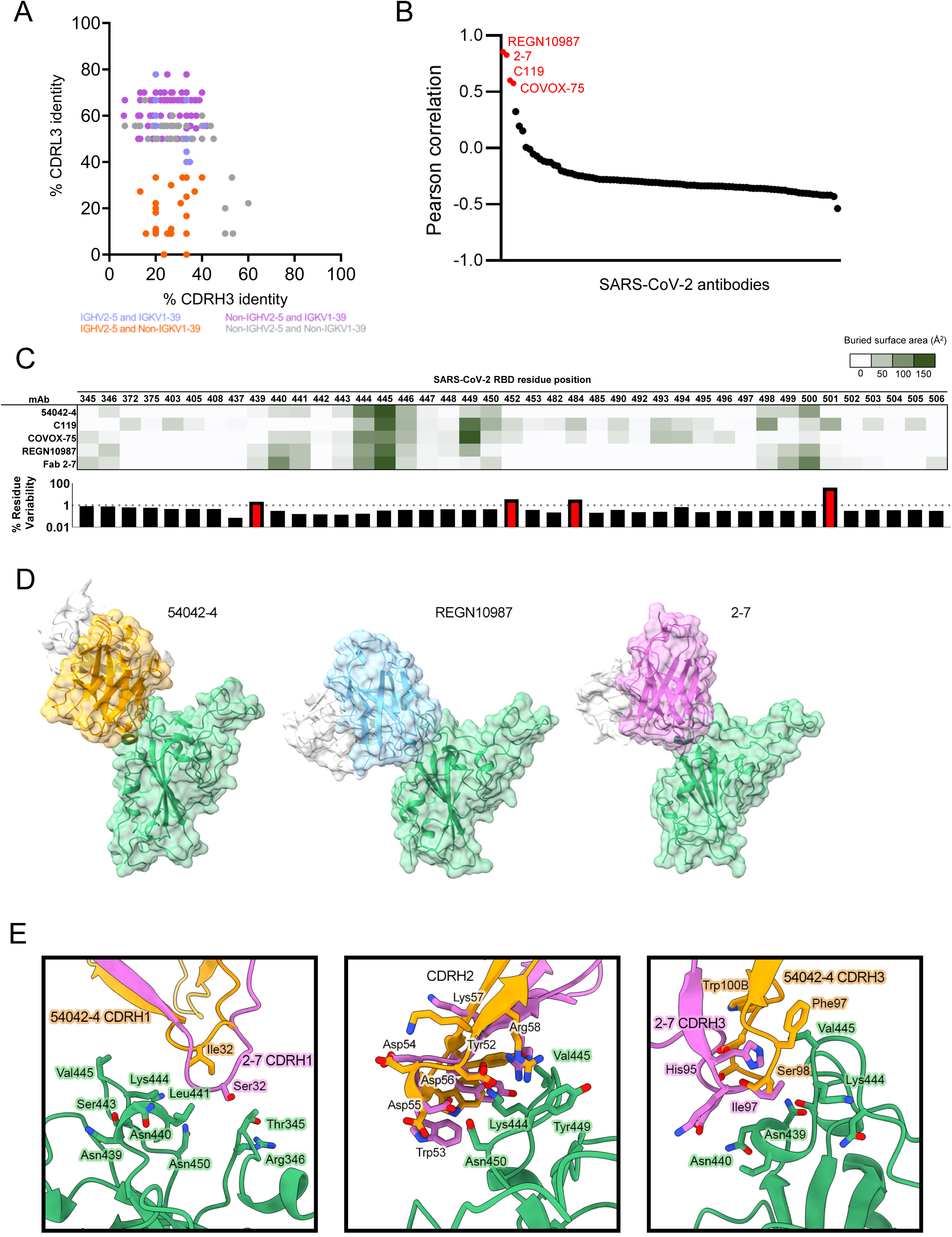
Sequence and structural comparison of 54042-4 to known SARS-CoV-2 antibodies. **(A)** Amino acid CDRH3 identity to 54042-4 (x-axis) is plotted against CDRL3 identity to 54042-4 (y-axis) for paired heavy and light chain sequences obtained from the CoV-AbDab database. Antibodies using the same heavy and light chain germline gene as 54042-4 (*IGHV2-5* and *IGKV1-39*) are shown in light blue. Antibodies using the *IGHV2-5* heavy chain gene and a non-*IGKV1-39* light chain gene are shown in orange. Additionally, antibodies using a non-*IGHV2-5* heavy chain gene and the *IGKV1-39* light chain gene, with CDRH3 or CDRL3 identity to 54042-4 of at least 50%, are highlighted in purple. Finally, antibodies that do not use *IGHV2-5* or *IGKV1-39*, but that have at least 50% identity to CDRH3 or CDRL3 of 54042-4, are shown in grey. **(B)** Pearson correlation of epitopes of known SARS-CoV-2 antibodies in comparison to 54042-4 antibody, with the four antibodies showing a statistically significant (p<0.05) positive correlation highlighted in red. **(C)** Heatmap (top) depicting buried surface area (Å^2^) at the SARS-CoV-2 RBD interface for the four antibodies with highest epitope correlations with 54042-4. Bar graph (bottom) showing the frequency (%) of substitutions at each given residue position in log scale, with a dashed line at 1% and residue positions with a frequency greater than 1% highlighted in red. **(D)** Distinct angles of approach of antibodies 54042-4 (heavy chain: orange, light chain: white), REGN10987 (heavy chain: blue, light chain: white) (PDB id: 6XDG), and 2-7 (heavy chain: pink, light chain: white) (PDB id: 7LSS) to the SARS-CoV-2 RBD (green). **(E)** Structural comparison of CDRH1, 2, and 3 of antibodies 54042-4 and 2-7. CDRH1 of 2-7 extends further than 54042-4, forming additional contacts with Thr345 and Arg346 of the RBD (left). The CDRH2 region of 2-7 approaches at a different angle, but maintains RBD contacts via Asp56 and Arg58 (center). The CDRH3 contacts of 2-7 and 54042-4 are divergent, with unique CDRH3 residues and RBD interactions (right).

Next, we compared the 54042-4 epitope to the epitopes of other known SARS-CoV-2 antibodies by computing pairwise correlations between the antibody-antigen buried surface areas for 54042-4 against a set of available SARS-CoV-2 antibody-antigen structures. The results revealed significant positive correlations with only four other antibodies: REGN10987(Hansen et al., 2020), 2-7 (Liu et al., 2020), C119 (Barnes et al., 2020), and COVOX-75 (Dejnirattisai et al., 2021) (**Figure 4B**). However, of these four antibodies, COVOX-75 has been reported as not a potent neutralizer (Dejnirattisai et al., 2021). C119 makes substantial contact with residues Asn501 and Glu484, indicating potential susceptibility of this antibody to substitutions at those positions that are currently associated with relatively high substitution rates (**Figure 4C**) and are present in several circulating SARS-CoV-2 VOCs (Alpert et al., 2021; Tegally et al., 2021). Further, both C119 and COVOX-75 have substantial buried surface area interactions with a number of additional residues compared to those in the epitope of 54042-4 (**Figure 4C**), suggesting that these two antibodies would be susceptible to a greater number of potential spike substitutions than 54042-4.

We also observed that while the epitopes of antibodies 2-7 and REGN10987 correlate well with that of 54042-4, these antibodies have distinct angles of antigen approach (**Figure 4D**). To quantify this observation, we aligned the RBDs from the 2-7 and REGN10987 complex structures with the RBD from the 54042-4 structure. Using the antibody coordinates when the respective RBDs were aligned, we computed the root mean square deviations (RMSD) between the C_α_ atoms in the FWR1-FWR3 regions of the antibody heavy and light chains. This resulted in RMSDs of 16.4 Å and 22 Å between 54042-4 versus 2-7 and REGN10987, respectively, confirming the substantial differences in the structural mode of antigen recognition by 54042-4 compared to 2-7 and REGN10987. Further, although 54042-4 and 2-7 both originate from the same *IGHV2-5* germline gene and share analogous RBD contacts in the CDRH2 region, these antibodies exhibit different CDRH1 and CDRH3 interactions (**Figure 4E**) and use a different light chain germline gene (*VK1-39* for 54042-4, and *VL2-14* for 2-7). Notably, both 2-7 and REGN10987 have greater interactions with RBD residues 439–441 compared to 54042-4, with buried surface areas of 172, 127, and 60 Å^2^ for 2-7, REGN10987, and 54042-4, respectively (**Figure 4C**), suggesting 2-7 and REGN10987 may be more prone to viral escape in that region. Indeed, the N439K substitution is present in several independent SARS-CoV-2 lineages and has been found to affect binding and neutralization by REGN10987 (Thomson et al., 2021).

Together, these data suggest that antibody 54042-4 utilizes a unique genetic signature and structural mode of antigen recognition that are distinct from other known SARS-CoV-2 antibodies.

### Antibody 54042-4 is not affected by current SARS-CoV-2 VOC substitutions

To identify substitutions capable of disrupting binding to antibody 54042-4, we next performed shotgun alanine-scanning mutagenesis of the SARS-CoV-2 RBD (Davidson, 2014). The only substitutions tested that substantially affected binding in comparison to an RBD antibody control were K444A, V445A, G446A, and P499A (**Figure 5A)**, which all fall within the 54042-4 epitope (**Figure 3C, D, and Supplemental Figure 3A**). Next, to assess the functional effect of substitutions within the 54042-4 epitope, we tested neutralization against VSV-SARS-CoV-2 chimeras containing substitutions at K444R/T/E/N, G446D, or Q498R. These specific substitutions were chosen based on their generation from neutralization-escape experiments with monotreatment at saturating concentration of antibodies COV2-2130 (shown to compete with 54042-4, **Figure 2D**) and COV2-2499 (a known COV2-2130 competitor) (Greaney et al, 2021).These experiments revealed that the chimeric VSVs with substitutions at Lys444, Gly446, and Gln498 were resistant to neutralization by 54042-4 (**Figure 5B**). Together, the alanine-scanning and neutralization-escape experiments indicated that 54042-4 recognition of spike may be sensitive to substitutions at residues K444, V445, G446, Q498, and P499. However, analysis of currently circulating SARS-CoV-2 isolates from the GISAID database as of May 6, 2021 (Elbe, 2017) revealed that substitutions at these five residue positions are only present at low levels (**Figure 5C**). Further, virtually all of the 54042-4 epitope residues (**Supplemental Figure 3A**) are highly conserved in circulating SARS-CoV-2 lineages (**Figure 5C**). The only exception is residue N439, which has a substitution frequency of 2.1% (**Figure 5C**); however, this residue makes only minimal contacts with antibody 54042-4 (**Supplemental Figure 3A**), suggesting that residue N439 may not be critical for 54042-4 recognition of the SARS-CoV-2 spike.

**Figure 5:**
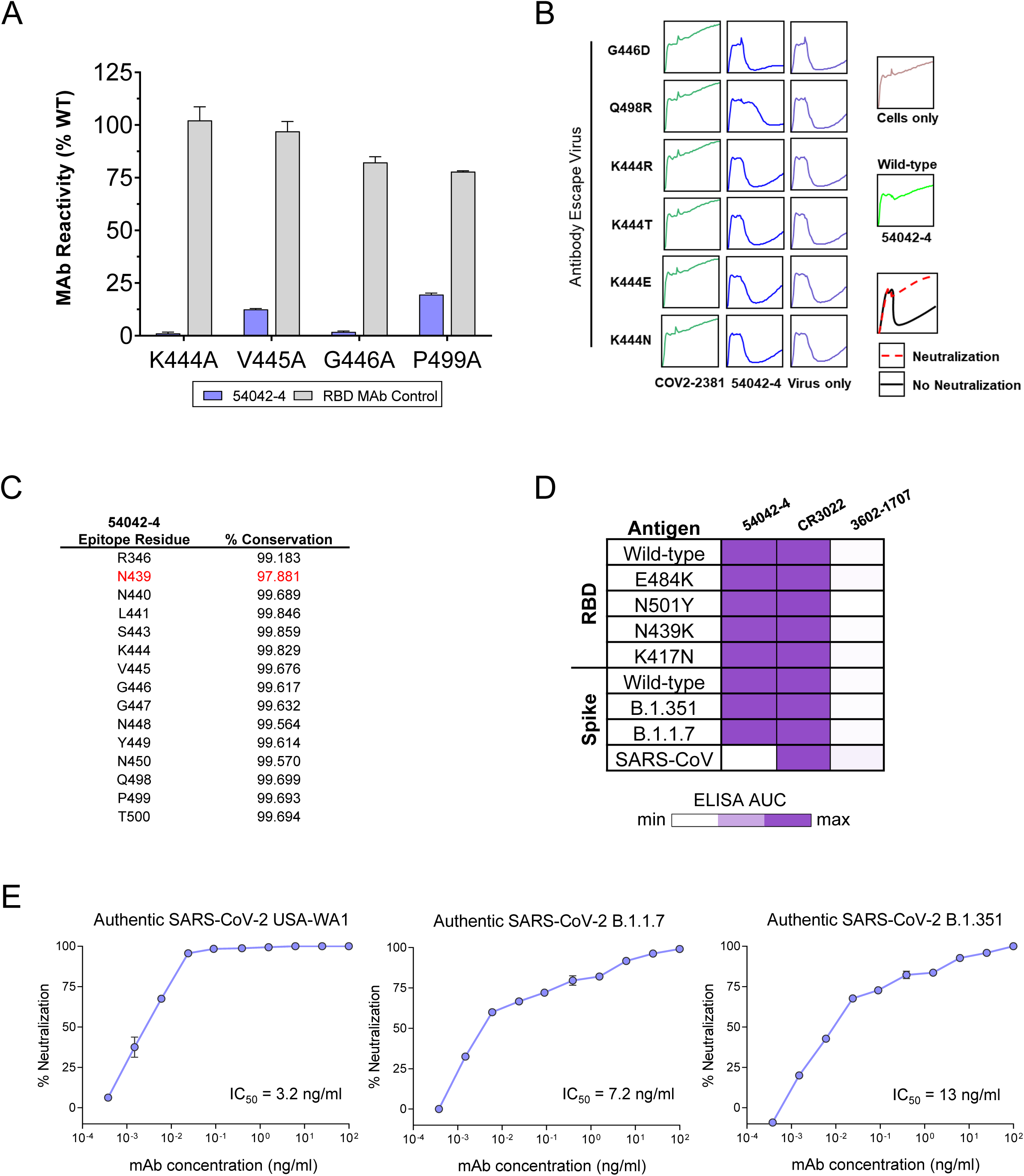
Functional characterization of antibody 54042-4. **(A)** Binding data of 54042-4 antibody to a shotgun alanine mutagenesis screening library of the SARS-CoV-2 RBD (Wuhan-Hu-1 strain). Residues displayed are the alanine mutations that resulted in the biggest loss of binding to 54042-4 yet still retained signal with the RBD antibody control. **(B)** RTCA Neutralization of VSV SARS-CoV-2 chimera variants harboring specific mutations. Cell sensograms are shown in boxes corresponding to mutations indicated in each row. Columns (from left to right) are each chimera treated with antibody2381 and 54042-4 and virus only control. Neutralization of 54042-4 of USA-WA1 strain and cells only are indicated on the right. COV2-2381 was chosen as a positive control due to its distinct epitope footprint from the selected mutations. **(C)** 54042-4 epitope residues (non-zero buried surface area on SARS-CoV-2 RBD) with their associated % conservation (the percentage of deposited sequences containing the highest-frequency amino acid at that position) in the GISAID database. The only 54042-4 epitope residue with a % conservation of less than 99%, N439, is highlighted in red. **(D)** ELISA AUC of 54042-4, CR3022, and an influenza HA-specific negative control antibody 3602-1707. AUC is displayed as a heatmap with a value of 0 corresponding to white, 50% maximum as light-purple, and maximum AUC as purple. **(E)** Authentic SARS-CoV-2 neutralization of USA-WA1, B.1.1.7, and B.1.351 is depicted as a function of antibody concentration.

To investigate the ability of antibody 54042-4 to recognize current SARS-CoV-2 VOCs, we next performed ELISAs to test binding of 54042-4 to RBD proteins containing substitutions found in one or more VOCs. These substitutions included K417N found in many isolates in the B.1.351 lineage, as well as E484K (B.1.351, P.1), N501Y (B.1.1.7, B.1.351, P.1), as well as N439K found in lineages B.1.141 and B.1.258 (Thomson et al., 2021). Notably, antibody 54042-4 bound to these RBD variants at a similar level compared to the binding to the RBD from the Wuhan-1 isolate (**Figure 5D, Supplemental Figure 4A**). These results are consistent with the structural observations that 54042-4 makes only minimal contacts with residue 439, and that none of the other RBD substitutions were at residues in the 54042-4 epitope (**Supplemental Figure 3A**). Binding of antibody 54042-4 also was not affected in the context of S ECD proteins that included deletions and substitutions in the S1 domain of the B.1.351 and B.1.1.7 VOCs (**Figure 5D, Supplemental Figure 4B,C**). Importantly, we also tested the ability of 54042-4 to neutralize authentic SARS-CoV-2 USA-WA1, B.1.1.7, and B.1.351 SARS-CoV-2 variants. Consistent with the ELISA data, 54042-4 neutralized each virus potently with IC50s of 3.2, 7.2, and 13 ng/mL, respectively (**Figure 5E**). Together, these data indicate that 54042-4 can be an effective countermeasure against currently circulating SARS-CoV-2 variants.

## Discussion

SARS-CoV-2 neutralizing antibody discovery efforts have produced an extensive panel of antibodies that show a wide range of functional effects, yet most antibodies discovered to date cluster into several classes based on RBD-binding orientation, ACE2 antagonism, and cross-reactivity to related SARS-like coronaviruses (Greaney et al, 2021). Here, we report the identification of 54042-4, an antibody that exhibited ultra-potent SARS-CoV-2 neutralization against USA-WA1 as well as the currently circulating VOCs B.1.1.7 and B.1.351. While the epitope of antibody 54042-4 showed partial overlap with that of several other known RBD antibodies, our findings revealed an overall unique mode of SARS-CoV-2 spike recognition, paired with an uncommon genetic signature that distinguishes 54042-4 from other SARS-CoV-2 antibodies. Notably, important differences were observed even for the four antibodies with the highest epitope correlations to 54042-4, with all four of these antibodies exhibiting substantially greater contacts with one or more known residues associated with currently circulating VOCs (**Figure 4C**). The discovery of antibody 54042-4 is therefore a unique addition to the limited set of antibodies with a high potential for effectively counteracting current SARS-CoV-2 VOCs.

The increased spread of several SARS-CoV-2 VOCs over the past few months has emphasized the need for continued surveillance of vaccine efficacy against the evolving virus targets. The increased transmission rates of the B.1.1.7 lineage are likely a product of enhanced ACE2 affinity for the SARS-CoV-2 RBD (Starr et al., 2020), and not a result of escape from pre-existing antibodies in convalescent or vaccinated individuals (Wang et al., 2021; Xie et al., 2021). Variants that encode the E484K substitution appear to pose a significantly higher risk of neutralization escape in vaccine recipients and individuals who have recovered from COVID-19 (Wang et al., 2021). Indeed, the rise of cases associated with the P.1 variant that harbors the E484K substitution (among others) in Manaus, Brazil is on a dangerous trajectory, despite having a 76% population seropositivity rate dating back to March 2020 (Sabino et al., 2021). In the context of vaccination, early vaccine trial data for Novavax against the B.1.351 lineage in South Africa (also encoding the E484K substitution) demonstrated a significant decrease in efficacy (Wadman, 2021). These observations underscore the ongoing need for genomic surveillance to monitor the emergence and spread of new SARS-CoV-2 variants and their effects on population immunity.

In addition to vaccines, antibody therapeutics can play an important role for treating SARS-CoV-2 infections. Given the unknown future trajectory of the pandemic and the potential for emergence of VOCs that may escape neutralization by vaccine-elicited immunity, the development of a wide array of candidate antibody therapeutics that are insensitive to substitutions found in major VOCs may prove critical in the fight against COVID-19. However, current VOCs have already shown an ability to escape neutralization by a number of antibodies in clinical development (Chen et al., 2021; Wang et al., 2021). In contrast, our binding, neutralization, and structural data suggest that antibody 54042-4 is capable of avoiding all of the current major substitutions in circulating VOCs. Combined with those observations, the unique features of 54042-4 in comparison to other SARS-CoV-2 antibodies motivate further clinical development of this antibody to complement the existing pool of therapeutic countermeasures. As SARS-CoV-2 virus evolution continues due to various factors, such as a lack of vaccine access and the associated delayed vaccine rollout to underserved parts of the world, new VOCs are likely to keep emerging, with the potential to decrease or even abrogate protection induced by current vaccines. Antibody therapeutic development, especially focusing on broad protection against diverse SARS-CoV-2 variants, is therefore of continued significance.

## METHODS

### Donor Information

The donor had previous laboratory-confirmed COVID-19, 3 months prior to blood collection. The studies were reviewed and approved by the Institutional Review Board of Vanderbilt University Medical Center. The sample was obtained after written informed consent was obtained.

### Antigen Purification

A variety of recombinant soluble protein antigens were used in the LIBRA-seq experiment and other experimental assays.

Plasmids encoding residues 1–1208 of the SARS-CoV-2 spike with a mutated S1/S2 cleavage site, proline substitutions at positions 817, 892, 899, 942, 986 and 987, and a C-terminal T4-fibritin trimerization motif, an 8x HisTag, and a TwinStrepTag (SARS-CoV-2 spike HP); 1–1208 of the SARS-CoV-2 spike with a mutated S1/S2 cleavage site, proline substitutions at positions 817, 892, 899, 942, 986 and 987, a glycine mutation at 614, and a C-terminal T4-fibritin trimerization motif, an 8x HisTag, and a TwinStrepTag (SARS-CoV-2 spike HP D614G) 1–1208 of the SARS-CoV-2 spike with a mutated S1/S2 cleavage site, proline substitutions at positions 817, 892, 899, 942, 986 and 987, as well as mutations L18F, D80A, L242-244L del, R246I, K417N, E484K, N501Y, and a C-terminal T4-fibritin trimerization motif, an 8x HisTag, and a TwinStrepTag (SARS-CoV-2 spike HP B.1.351); 1–1208 of the SARS-CoV-2 spike with a mutated S1/S2 cleavage site, proline substitutions at positions 817, 892, 899, 942, 986 and 987, as well as mutations 69-70del, Y144del, N501Y, A570D, P681H, and a C-terminal T4-fibritin trimerization motif, an 8x HisTag, and a TwinStrepTag (SARS-CoV-2 spike HP B.1.1.7); residues 1-1190 of the SARS-CoV spike with proline substitutions at positions 968 and 969, and a C-terminal T4-fibritin trimerization motif, an 8x HisTag, and a TwinStrepTag (SARS-CoV S-2P); residues 1-1291 of the MERS-CoV spike with a mutated S1/S2 cleavage site, proline substitutions at positions 1060 and 1061, and a C-terminal T4-fibritin trimerization motif, an AviTag, an 8x HisTag, and a TwinStrepTag (MERS-CoV S-2P Avi); residues 1-1278 of the HCoV-OC43 spike with proline substitutions at positions 1070 and 1071, and a C-terminal T4-fibritin trimerization motif, an 8x HisTag, and a TwinStrepTag (HCoV-OC43 S-2P); residues 319–591 of SARS-CoV-2 S with a C-terminal monomeric human IgG Fc-tag and an 8x HisTag (SARS-CoV-2 RBD-SD1); residues 367–589 of MERS-CoV S with a C-terminal monomeric human IgG Fc-tag and an 8x HisTag (MERS-CoV RBD); residues 306–577 of MERS-CoV S with a C-terminal monomeric human IgG Fc-tag and an 8x HisTag (SARS-CoV RBD-SD1) were transiently transfected into FreeStyle293F cells (Thermo Fisher) using polyethylenimine. For all antigens with the exception of SARS-CoV-2 S HP, cells were treated with 1 µM kifunensine to ensure uniform glycosylation three hours post-transfection. Transfected supernatants were harvested after 6 days of expression. SARS-CoV-2 RBD-SD1 was purified using Protein A resin (Pierce), SARS-CoV-2 S HP, MERS-CoV S-2P, and HCoV-OC43 S-2P were purified using StrepTactin resin (IBA). Affinity-purified SARS-CoV-2 RBD-SD1 was further purified over a Superdex75 column (GE Life Sciences). SARS-CoV-2 S HP, SARS-CoV-2 S HP B.1.351, SARS-CoV-2spikeHP B.1.1.7, SARS-CoV S-2P, MERS-CoV S-2P, HCoV-HKU1 S-2P and HCoV-OC43 S-2P were purified over a Superose6 Increase column (GE Life Sciences). HCoV-NL63 and HCoV-229E alpha coronavirus spike proteins were purchased from Sino Biological. SARS-CoV-2 S1, SARS-CoV-2 S2, and SARS-CoV-2 NTD truncated proteins were purchased from the commercial vendor, Sino Biological.

For the HIV-1 gp140 SOSIP variant from strain ZM197 (clade C) recombinant, soluble antigens contained an AviTag and were expressed in Expi293F cells using polyethylenimine transfection reagent and cultured. FreeStyle F17 expression medium supplemented with pluronic acid and glutamine was used. The cells were cultured at 37°C with 8% CO_2_ saturation and shaking. After 5-7 days, cultures were centrifuged and supernatant was filtered and run over an affinity column of agarose bound *Galanthus nivalis* lectin. The column was washed with PBS and antigens were eluted with 30 mL of 1M methyl-a-D-mannopyranoside. Protein elutions were buffer exchanged into PBS, concentrated, and run on a Superdex 200 Increase 10/300 GL Sizing column on the AKTA FPLC system. Fractions corresponding to correctly folded protein were collected, analyzed by SDS-PAGE and antigenicity was characterized by ELISA using known monoclonal antibodies specific to each antigen. Avitagged antigens were biotinylated using BirA biotin ligase (Avidity LLC).

Recombinant NC99 HA protein consists of the HA ectodomain with a point mutation at the sialic acid-binding site (Y98F) to abolish non-specific interactions, a T4 fibritin foldon trimerization domain, AviTag, and hexahistidine-tag, and were expressed in Expi 293F mammalian cells using Expifectamine 293 transfection reagent (Thermo Fisher Scientific) cultured for 4-5 days. Culture supernatant was harvested and cleared as above, and then adjusted pH and NaCl concentration by adding 1M Tris-HCl (pH 7.5) and 5M NaCl to 50 mM and 500 mM, respectively. Ni Sepharose excel resin (GE Healthcare) was added to the supernatant to capture hexahistidine tag. Resin was separated on a column by gravity and captured HA protein was eluted by a Tris-NaCl (pH 7.5) buffer containing 300 mM imidazole. The eluate was further purified by a size exclusion chromatography with a HiLoad 16/60 Superdex 200 column (GE Healthcare). Fractions containing HA were concentrated, analyzed by SDS-PAGE and tested for antigenicity by ELISA with known antibodies.

Spike protein used for cryo-EM was expressed by transiently transfecting plasmid encoding the HexaPro spike variant (Hsieh et al., 2020) containing additional S383C and D985C substitutions (Henderson et al., 2020) with a C-terminal TwinStrep tag into FreeStyle 293-F cells (Thermo Fisher) using polyethyleneimine. 5 μM kifunensine was added 3h post-transfection. The cell culture was harvested four days after transfection and the spike-containing medium was separated from the cells by centrifugation. Supernatants were passed through a 0.22 µm filter and passaged over StrepTactin resin (IBA). Further purification was achieved by size-exclusion chromatography using a Superose 6 10/300 column (GE Healthcare) in buffer containing 2 mM Tris pH 8.0, 200 mM NaCl and 0.02% NaN3.

### DNA-barcoding of Antigens

We used oligos that possess 15 bp antigen barcode, a sequence capable of annealing to the template switch oligo that is part of the 10X bead-delivered oligos, and contain truncated TruSeq small RNA read 1 sequences in the following structure: 5’-CCTTGGCACCCGAGAATTCCANNNNNNNNNNNNNCCCATATAAGA*A*A-3’, where Ns represent the antigen barcode. We used the following antigen barcodes: We used the following antigen barcodes: GCAGCGTATAAGTCA (SARS-CoV-2 S), AACCCACCGTTGTTA (SARS-CoV-2 S D614G), GCTCCTTTACACGTA (SARS-CoV S), GGTAGCCCTAGAGTA (MERS-CoV S), AGACTAATAGCTGAC (HCoV-OC43 S), GACAAGTGATCTGCA (HCoV-NL63 S), GTGTGTTGTCCTATG (HCoV-229E S), TACGCCTATAACTTG (ZM197 EnV), TCATTTCCTCCGATT (HA NC99), TGGTAACGACAGTCC (SARS-CoV RBD-SD1), TTTCAACGCCCTTTC (SARS-CoV-2 RBD-SD1), GTAAGACGCCTATGC (MERS-CoV RBD), CAGTAAGTTCGGGAC(SARS-CoV-2 NTD), Oligos were ordered from IDT with a 5’ amino modification and HPLC purified.

For each antigen, a unique DNA barcode was directly conjugated to the antigen itself. In particular, 5’amino-oligonucleotides were conjugated directly to each antigen using the Solulink Protein-Oligonucleotide Conjugation Kit (TriLink cat no. S-9011) according to manufacturer’s instructions. Briefly, the oligo and protein were desalted, and then the amino-oligo was modified with the 4FB crosslinker, and the biotinylated antigen protein was modified with S-HyNic. Then, the 4FB-oligo and the HyNic-antigen were mixed together. This causes a stable bond to form between the protein and the oligonucleotide. The concentration of the antigen-oligo conjugates was determined by a BCA assay, and the HyNic molar substitution ratio of the antigen-oligo conjugates was analyzed using the NanoDrop according to the Solulink protocol guidelines. AKTA FPLC was used to remove excess oligonucleotide from the protein-oligo conjugates, which were also verified using SDS-PAGE with a silver stain. Antigen-oligo conjugates were also used in flow cytometry titration experiments.

### Antigen-specific B cell sorting

Cells were stained and mixed with DNA-barcoded antigens and other antibodies, and then sorted using fluorescence activated cell sorting (FACS). First, cells were counted and viability was assessed using Trypan Blue. Then, cells were washed three times with DPBS supplemented with 0.1% Bovine serum albumin (BSA). Cells were resuspended in DPBS-BSA and stained with cell markers including viability dye (Ghost Red 780), CD14-APC-Cy7, CD3-FITC, CD19-BV711, and IgG-PE-Cy5. Additionally, antigen-oligo conjugates were added to the stain. After staining in the dark for 30 minutes at room temperature, cells were washed three times with DPBS-BSA at 300 g for five minutes. Cells were then incubated for 15 minutes at room temperature with Streptavidin-PE to label cells with bound antigen. Cells were washed three times with DPBS-BSA, resuspended in DPBS, and sorted by FACS. Antigen positive cells were bulk sorted and delivered to the Vanderbilt Technologies for Advanced Genomics (VANTAGE) sequencing core at an appropriate target concentration for 10X Genomics library preparation and subsequent sequencing. FACS data were analyzed using FlowJo.

### Sample preparation, library preparation, and sequencing

Single-cell suspensions were loaded onto the Chromium Controller microfluidics device (10X Genomics) and processed using the B-cell Single Cell V(D)J solution according to manufacturer’s suggestions for a target capture of 10,000 B cells per 1/8 10X cassette, with minor modifications in order to intercept, amplify and purify the antigen barcode libraries as previously described (Setliff et al., 2019).

### Sequence processing and bioinformatics analysis

We utilized and modified our previously described pipeline to use paired-end FASTQ files of oligo libraries as input, process and annotate reads for cell barcode, unique molecular identifier (UMI), and antigen barcode, and generate a cell barcode - antigen barcode UMI count matrix (Setliff et al., 2019). BCR contigs were processed using Cell Ranger (10X Genomics) using GRCh38 as reference. Antigen barcode libraries were also processed using Cell Ranger (10X Genomics). The overlapping cell barcodes between the two libraries were used as the basis of the subsequent analysis. We removed cell barcodes that had only non-functional heavy chain sequences as well as cells with multiple functional heavy chain sequences and/or multiple functional light chain sequences, reasoning that these may be multiplets. Additionally, we aligned the BCR contigs (filtered_contigs.fasta file output by Cell Ranger, 10X Genomics) to IMGT reference genes using HighV-Quest (Alamyar et al., 2012). The output of HighV-Quest was parsed using ChangeO(Gupta et al., 2015) and merged with an antigen barcode UMI count matrix. Finally, we determined the LIBRA-seq score for each antigen in the library for every cell as previously described(Setliff et al., 2019).

### Antibody expression and purification

For each antibody, variable genes were inserted into custom plasmids encoding the constant region for the IgG1 heavy chain as well as respective lambda and kappa light chains (pTwist CMV BetaGlobin WPRE Neo vector, Twist Bioscience). Antibodies were expressed in Expi293F mammalian cells (Thermo Fisher Scientific) by co-transfecting heavy chain and light chain expressing plasmids using polyethylenimine transfection reagent and cultured for 5-7 days. Cells were maintained in FreeStyle F17 expression medium supplemented at final concentrations of 0.1% Pluronic Acid F-68 and 20% 4mM L-Glutamine. These cells were cultured at 37°C with 8% CO_2_ saturation and shaking. After transfection and 5-7 days of culture, cell cultures were centrifuged and supernatant was 0.45 μm filtered with Nalgene Rapid Flow Disposable Filter Units with PES membrane. Filtered supernatant was run over a column containing Protein A agarose resin equilibrated with PBS. The column was washed with PBS, and then antibodies were eluted with 100 mM Glycine HCl at 2.7 pH directly into a 1:10 volume of 1M Tris-HCl pH 8.0. Eluted antibodies were buffer exchanged into PBS 3 times using Amicon Ultra centrifugal filter units and concentrated. Antibodies were analyzed by SDS-PAGE.

### High-throughput antibody expression

For high-throughput production of recombinant antibodies, approaches were used that are designated as microscale. For antibody expression, microscale transfection were performed (∼1 ml per antibody) of CHO cell cultures using the Gibco ExpiCHO Expression System and a protocol for deep 96-well blocks (Thermo Fisher Scientific). In brief, synthesized antibody-encoding DNA (∼2 μg per transfection) was added to OptiPro serum free medium (OptiPro SFM), incubated with ExpiFectamine CHO Reagent and added to 800 µl of ExpiCHO cell cultures into 96-deep-well blocks using a ViaFlo 384 liquid handler (Integra Biosciences). The plates were incubated on an orbital shaker at 1,000 r.p.m. with an orbital diameter of 3 mm at 37 °C in 8% CO_2_. The next day after transfection, ExpiFectamine CHO Enhancer and ExpiCHO Feed reagents (Thermo Fisher Scientific) were added to the cells, followed by 4 d incubation for a total of 5 d at 37 °C in 8% CO_2_. Culture supernatants were collected after centrifuging the blocks at 450*g* for 5 min and were stored at 4°C until use. For high-throughput microscale antibody purification, fritted deep-well plates were used containing 25 μl of settled protein G resin (GE Healthcare Life Sciences) per well. Clarified culture supernatants were incubated with protein G resin for antibody capturing, washed with PBS using a 96-well plate manifold base (Qiagen) connected to the vacuum and eluted into 96-well PCR plates using 86 μl of 0.1 M glycine-HCL buffer pH 2.7. After neutralization with 14 μl of 1 M Tris-HCl pH 8.0, purified antibodies were buffer-exchanged into PBS using Zeba Spin Desalting Plates (Thermo Fisher Scientific) and stored at 4°C until use.

### ELISA

To assess antibody binding, soluble protein was plated at 2 μg/ml overnight at 4°C. The next day, plates were washed three times with PBS supplemented with 0.05% Tween-20 (PBS-T) and coated with 5% milk powder in PBS-T. Plates were incubated for one hour at room temperature and then washed three times with PBS-T. Primary antibodies were diluted in 1% milk in PBS-T, starting at 10 μg/ml with a serial 1:5 dilution and then added to the plate. The plates were incubated at room temperature for one hour and then washed three times in PBS-T. The secondary antibody, goat anti-human IgG conjugated to peroxidase, was added at 1:10,000 dilution in 1% milk in PBS-T to the plates, which were incubated for one hour at room temperature. Plates were washed three times with PBS-T and then developed by adding 3,3′,5,5′-tetramethylbenzidine (TMB) substrate to each well. The plates were incubated at room temperature for ten minutes, and then 1N sulfuric acid was added to stop the reaction. Plates were read at 450 nm. Data are represented as mean ± SEM for one ELISA experiment. ELISAs were repeated 2 or more times. The area under the curve (AUC) was calculated using GraphPad Prism 9.0.1.

### Competition ELISA

Competition ELISA was performed as done previously (Zost et al., 2020). Briefly, antibodies were biotinylated using NHS-PEG4-biotin (Thermo Fisher Scientific, cat# A39259) according to manufacturer protocol. Following biotinylation, specific binding of biotinylated antibodies was confirmed using ELISA. Wells of 384-well microtiter plates were coated with 1 µg/mL SARS-CoV-2 S HP protein at 4°C overnight. Plates were washed with PBS-T and blocked for 1 h with blocking buffer (1% BSA in PBS-T). Plates were then washed with PBS-T and unlabeled antibodies were added at a concentration of 10 µg/mL in a total volume of 25 µL blocking buffer and incubated 1 h. Without washing, biotinylated antibodies diluted in blocking buffer were added directly to each well in a volume of 5 µL per well (such that the final concentrations of each biotinylated antibody were equal to the respective EC_90_ of each antibody), and then incubated for 30 min at ambient temperature. Plates were then washed with PBS-T and incubated for 1 h with HRP-conjugated avidin (Sigma, 25 µL of a 1:3,500 dilution in blocking buffer). Plates were washed with PBS-T and 25 µL TMB substrate was added to each well. After sufficient development, the reactions were quenched by addition 25 µL 1M HCl and absorbance at 450 nm was quantified using a plate reader. After subtracting the background signal, the signal obtained for binding of the biotin-labeled reference antibody in the presence of the unlabeled tested antibody was expressed as a percentage of the binding of the reference antibody in the presence of 10 µg/mL of the anti-dengue antibody DENV 2D22, which served as a no-competition control. Tested antibodies were considered competing if their presence reduced the reference antibody binding by more than 60% and non-competing if the signal was reduced by less than 30%.

### Real-time Cell Analysis (RTCA) Neutralization Assay Screen

To screen for neutralizing activity in the panel of recombinantly expressed antibodies, we used a high-throughput and quantitative RTCA assay and xCelligence RTCA HT Analyzer (ACEA Biosciences) that assesses kinetic changes in cell physiology, including virus-induced cytopathic effect (CPE). Twenty µl of cell culture medium (DMEM supplemented with 2% FBS) was added to each well of a 384-well E-plate using a ViaFlo384 liquid handler (Integra Biosciences) to obtain background reading. Six thousand (6,000) Vero-furin cells in 20 μl of cell culture medium were seeded per well, and the plate was placed on the analyzer. Sensograms were visualized using RTCA HT software version 1.0.1 (ACEA Biosciences). For a screening neutralization assay, equal amounts of virus were mixed with micro-scale purified antibodies in a total volume of 40 μL using DMEM supplemented with 2% FBS as a diluent and incubated for 1 h at 37 °C in 5% CO2. At ∼17–20 h after seeding the cells, the virus–antibody mixtures were added to the cells in 384-well E-plates. Wells containing virus only (in the absence of antibody) and wells containing only Vero cells in medium were included as controls. Plates were measured every 8–12 h for 48–72 h to assess virus neutralization. Micro-scale antibodies were assessed in four 5-fold dilutions (starting from a 1:20 sample dilution), and their concentrations were not normalized. Neutralization was calculated as the percent of maximal cell index in control wells without virus minus cell index in control (virus-only) wells that exhibited maximal CPE at 40–48 h after applying virus–antibody mixture to the cells. An antibody was classified as fully neutralizing if it completely inhibited SARS-CoV-2-induced CPE at the highest tested concentration, while an antibody was classified as partially neutralizing if it delayed but did not fully prevent CPE at the highest tested concentration(Zost et al., 2020)).

### RTCA Potency Neutralization Screening Assay

To determine neutralizing activity of IgG, we used real-time cell analysis (RTCA) assay on an xCELLigence RTCA MP Analyzer (ACEA Biosciences Inc.) that measures virus-induced cytopathic effect (CPE) (Suryadevara N et al., 2021). Briefly, 50 μL of cell culture medium (DMEM supplemented with 2% FBS) was added to each well of a 96-well E-plate using a ViaFlo384 liquid handler (Integra Biosciences) to obtain background reading. A suspension of 18,000 Vero-E6 cells in 50 μL of cell culture medium was seeded in each well, and the plate was placed on the analyzer. Measurements were taken automatically every 15 min, and the sensograms were visualized using RTCA software version 2.1.0 (ACEA Biosciences Inc). VSV-SARS-CoV-2 (0.01 MOI, ∼120 PFU per well) was mixed 1:1 with a dilution of antibody in a total volume of 100 μL using DMEM supplemented with 2% FBS as a diluent and incubated for 1 h at 37°C in 5% CO2. At 16 h after seeding the cells, the virus-antibody mixtures were added in replicates to the cells in 96-well E-plates. Triplicate wells containing virus only (maximal CPE in the absence of antibody) and wells containing only Vero cells in medium (no-CPE wells) were included as controls. Plates were measured continuously (every 15 min) for 48 h to assess virus neutralization. Normalized cellular index (CI) values at the endpoint (48 h after incubation with the virus) were determined using the RTCA software version 2.1.0 (ACEA Biosciences Inc.). Results are expressed as percent neutralization in a presence of respective antibody relative to control wells with no CPE minus CI values from control wells with maximum CPE. RTCA IC50 values were determined by nonlinear regression analysis using Prism software.

### Epitope mapping of antibodies by alanine scanning

Epitope mapping was performed essentially as described previously (Davidson, 2014) using a SARS-CoV-2 (strain Wuhan-Hu-1) spike protein RBD shotgun mutagenesis mutation library, made using an expression construct for full-length spike protein. 184 residues of the RBD (between spike residues 335 and 526) were mutated individually to alanine, and alanine residues to serine and clones arrayed in 384-well plates, one mutant per well. Antibody binding to each mutant clone was determined, in duplicate, by high-throughput flow cytometry. Each spike protein mutant was transfected into HEK-293T cells and allowed to express for 22 hrs. Cells were fixed in 4% (v/v) paraformaldehyde (Electron Microscopy Sciences), and permeabilized with 0.1% (w/v) saponin (Sigma-Aldrich) in PBS plus calcium and magnesium (PBS++) before incubation with antibodies diluted in PBS++, 10% normal goat serum (Sigma), and 0.1% saponin. Antibody screening concentrations were determined using an independent immunofluorescence titration curve against cells expressing wild-type spike protein to ensure that signals were within the linear range of detection. Antibodies were detected using 3.75 μg/mL of AlexaFluor488-conjugated secondary antibody (Jackson ImmunoResearch Laboratories) in 10% normal goat serum with 0.1% saponin. Cells were washed three times with PBS++/0.1% saponin followed by two washes in PBS, and mean cellular fluorescence was detected using a high-throughput Intellicyte iQue flow cytometer (Sartorius). Antibody reactivity against each mutant spike protein clone was calculated relative to wild-type spike protein reactivity by subtracting the signal from mock-transfected controls and normalizing to the signal from wild-type S-transfected controls. Mutations within clones were identified as critical to antibody binding if they did not support reactivity of the test antibody, but supported reactivity of other SARS-CoV-2 antibodies. This counter-screen strategy facilitates the exclusion of spike mutants that are locally misfolded or have an expression defect.

### Plaque reduction neutralization test (PRNT)

The virus neutralization with live authentic SARS-CoV-2 virus (USA-WA1) was performed in the BSL-3 facility of the Galveston National Laboratory using Vero E6 cells (ATCC CRL-1586) following the standard procedure. Briefly, Vero E6 cells were cultured in 96-well plates (10^4^ cells/well). Next day, 4-fold serial dilutions of antibodies were made using MEM-2% FBS, as to get an initial concentration of 100 µg/ml. Equal volume of diluted antibodies (60 µl) were mixed gently with original SARS-CoV-2 or B.1.1.7 variant or B.1.351 variant (60 µl containing 200 pfu) and incubated for 1 h at 37°C/5% CO_2_ atmosphere. The virus-serum mixture (100 µl) was added to cell monolayer in duplicates and incubated for 1 at h 37°C/5% CO2 atmosphere. Later, virus-serum mixture was discarded gently, and cell monolayer was overlaid with 0.6% methylcellulose and incubated for 2 days. The overlay was removed, and the plates were fixed in 4% paraformaldehyde twice following BSL-3 protocol. The plates were stained with 1% crystal violet and virus-induced plaques were counted. The percent neutralization and/or NT_50_ of antibody was calculated by dividing the plaques counted at each dilution with plaques of virus-only control. For antibodies, the inhibitory concentration at 50% (IC_50_) values were calculated in GraphPad Prism software by plotting the midway point between the upper and lower plateaus of the neutralization curve among dilutions.

### BioLayer Interferometry (BLI)

Purified 54042-4 IgG was immobilized to AHC sensortips (FortéBio) to a response level of approximately 1.4 nm in a buffer composed of 10 mM HEPES pH 7.5, 150 mM NaCl, 3 mM EDTA, 0.05% Tween 20 and 0.1% (w/v) BSA. Immobilized IgG was then dipped into wells containing four-fold dilutions of SARS-CoV-2 RBD-SD1 ranging in concentration from 100-1.5625 nM, to measure association. Dissociation was measured by dipping sensortips into wells containing only running buffer. Data were reference subtracted and kinetics were calculated in Octet Data Analysis software v10.0 using a 1:1 binding model.

### ACE2 binding inhibition assay

96-well plates were coated with 2 μg/mL purified recombinant SARS-CoV-2 at 4°C overnight. The next day, plates were washed three times with PBS supplemented with 0.05% Tween-20 (PBS-T) and coated with 5% milk powder in PBS-T. Plates were incubated for one hour at room temperature and then washed three times with PBS-T. Purified antibodies were diluted in blocking buffer at 10 μg/mL in triplicate, added to the wells, and incubated at room temperature. Without washing, recombinant human ACE2 protein with a mouse Fc tag was added to wells for a final 0.4 μg/mL concentration of ACE2 and incubated for 40 minutes at room temperature. Plates were washed three times with PBS-T, and bound ACE2 was detected using HRP-conjugated anti-mouse Fc antibody and TMB substrate. The plates were incubated at room temperature for ten minutes, and then 1N sulfuric acid was added to stop the reaction. Plates were read at 450 nm. ACE2 binding without antibody served as a control. Experiment was done in biological replicate and technical triplicates.

### Neutralization escape

We used a real-time cell analysis assay (RTCA) and xCELLigence RTCA MP Analyzer (ACEA Biosciences Inc.) with modification of previously described assays (Gilchuk et al., 2020a; Weisblum et al., 2020, Suryadevara et al.,2021). Fifty (50) μL of cell culture medium (DMEM supplemented with 2% FBS) was added to each well of a 96-well E-plate to obtain a background reading. Eighteen thousand (18,000) Vero E6 cells in 50 μL of cell culture medium were seeded per each well, and plates were placed on the analyzer. Measurements were taken automatically every 15 min and the sensograms were visualized using RTCA software version 2.1.0 (ACEA Biosciences Inc). COV2-2130 or COV2-2499 or WT VSV-SARS-CoV-2 virus (5e3 plaque forming units [PFU] per well, ∼0.3 MOI) was mixed with a saturating neutralizing concentration of individual antibody (5 μg/mL) in a total volume of 100 μL and incubated for 1 h at 37°C. At 16-20 h after seeding the cells, the virus-antibody mixtures were added into 8 to 96 replicate wells of 96-well E-plates with cell monolayers. Wells containing only virus in the absence of antibody and wells containing only Vero E6 cells in medium were included on each plate as controls. Plates were measured continuously (every 15 min) for 72 h. The escapes from 54042-4 was confirmed by delayed CPE in wells containing antibody while mAb2381 was used as positive control.

### EM sample prep and data collection

To form the spike-Fab complex, a final concentration of 0.5 mg/mL spike protein and 5X molar excess of Fab were combined in buffer containing 2mM Tris-Cl pH 8.0, 200 mM NaCl, and 0.02% NaN_3_. The complex was incubated on ice for 30 min before 3 µL of the sample was deposited on Au-300 1.2/1.3 grids (UltrAuFoil) that had been plasma cleaned in a Solarus 950 plasma cleaner (Gatan) for 4 minutes using a 4:1 ratio of O_2_:H_2_. A force of -4 was used to blot excess liquid for 3 s using a Vitrobot Mark IV (Thermo Fisher) followed by plunge-freezing with liquid ethane. 3,762 micrographs were collected from a single grid using a Titan Krios (Thermo Fisher) equipped with a K3 detector (Gatan) with the stage set at a 30° tilt. SerialEM was used to collect movies at 29,000X nominal magnification with a calibrated pixel size of 0.81 Å/pixel. Additional details about data collection parameters can be found in **Supplemental Table 1.**

### Cryogenic electron microscopy (Cryo-EM)

Motion correction, CTF estimation, particle picking, and preliminary 2D classification were performed using cryoSPARC v3.2.0 live processing (Punjani et al., 2017). The final iteration of 2D class averaging distributed 374,669 particles into 60 classes using an uncertainty factor of 2. From that, 241,732 particles were used to perform an ab inito reconstruction with four classes followed by heterogeneous refinement of those four classes. Particles from the two highest quality classes were used for homogenous refinement of the best volume with applied C3 symmetry. Non-uniform refinement was performed on the resulting volume using per-particle defocus and per-group CTF optimizations applied (Punjani et al., 2020; Rubinstein and Brubaker, 2015). To improve the 54042-4 Fab-RBD density, C3 symmetry expansion was performed followed by local refinement using a mask created in ChimeraX that encompassed the entire 54042-4 Fab and RBD (Pettersen et al., 2021). Local refinement was performed using a pose/shift gaussian prior during alignment, 3° standard deviation of prior over rotation and 1 Å standard deviation of prior over shifts. Additionally, maximum alignment resolution was limited to 2.8 Å resolution to avoid over-refining. To improve map quality, the focused refinement volumes were processed using the DeepEMhancer(Sanchez-Garcia, 2021) tool via COSMIC^2^science gateway, which included the use of our refinement mask to help define noise while sharpening (Cianfrocco, 2017a; Cianfrocco, 2017b). An initial model was generated by docking PDBID: 6XKL (Hsieh et al., 2020) and a Fab model based on the 54042-4 sequence built using SAbPred ABodyBuilder (Dunbar et al., 2016) into map density via ChimeraX (Pettersen et al., 2021). The model was iteratively refined and completed using a combination of Phenix, Coot, and ISOLDE (Adams et al., 2002; Croll, 2018; Emsley and Cowtan, 2004). Details on structure validation and the full cryo-EM processing workflow can be found in **Supplemental Figures 5** and **6**.

### GISAID mutation frequency calculation

To evaluate the conservation of 54042-4 epitope residues, we utilized the GISAID database (Elbe, 2017) comprising sequences from 1229459 SARS-CoV-2 variants (as of May 6th, 2021). The spike glycoprotein sequences were extracted and translated, and pairwise sequence alignment with the reference sequence hCoV-19/Wuhan/WIV04/2019 was then performed. After removing incomplete sequences and sequences with alignment errors, the pairwise alignments for the remaining 1,148,887 spike protein sequences were combined to compute the conservation of each residue position using in-house perl scripts.

### RMSD calculation for the differences in angle of antigen approach for different antibodies

The SARS-CoV-2 spike receptor binding domain coordinates present in each antibody-antigen complex were aligned in PyMOL (The PyMOL Molecular Graphics System, Version 1.2r3pre, Schrödinger, LLC.) using an all-atom alignment with 5 cycles of outlier rejection of atom pairs having an RMSD greater than 2. The alignment was performed for RBD residues 329-529 in antibody 54042-4 (PDB ID: TBD chain A), 329-529 in antibody 2-7 (PDB ID: 7LSS chain B), and 333-526 in antibody REGN10987 (PDB ID: 6XDG chain A). This resulted in RMSD values of 0.751 Å between 54042-4 and REGN10987’s RBDs, 1.044 Å between 54042-4 and antibody 2-7’s RBDs, and 1.067 Å between REGN10987 and antibody 2-7’s RBDs with well-aligned epitope residues. Next, the residues comprising the N-termini through the end of framework region 3 were determined for the heavy and light chains of all three antibodies using IMGT Domain Gap Align (Alamyar et al., 2012). Each pair of antibodies was aligned using a pairwise sequence alignment of this region in PyMOL. Finally, the alpha carbon root mean square deviation between antibodies was calculated over this region in the heavy and light chains using residue pairs from the sequence alignment. RMSD values were calculated from 183, 183, and 180 alpha carbon pairs for the 54042-4 vs REGN1087, REGN1087 vs 2-7, and 54042-4 vs 2-7 comparisons respectively.

## QUANTIFICATION AND STATISTICAL ANALYSIS

ELISA error bars (standard error of the mean) were calculated using GraphPad Prism version 9.0.1.

**Supplemental Table 1:**
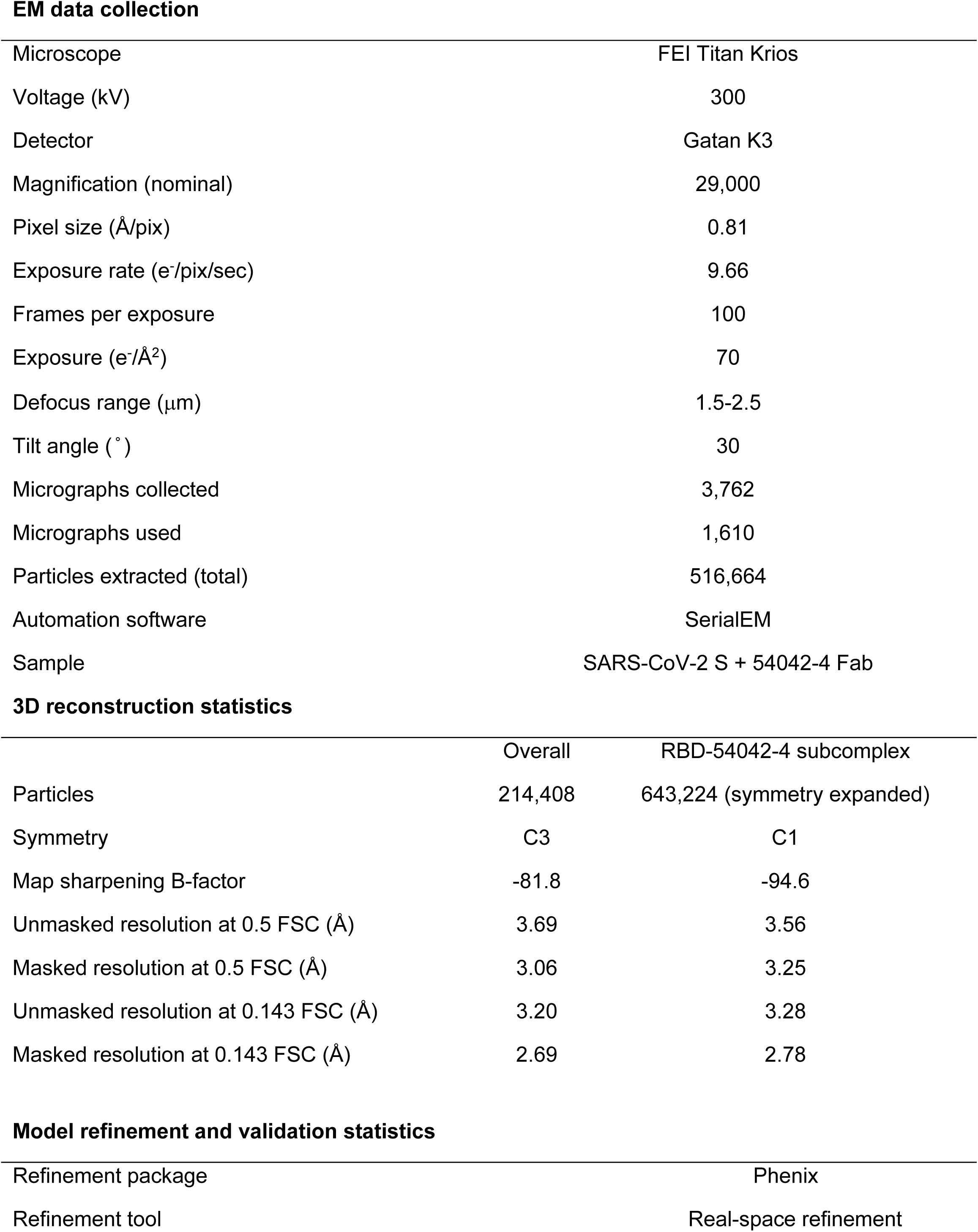
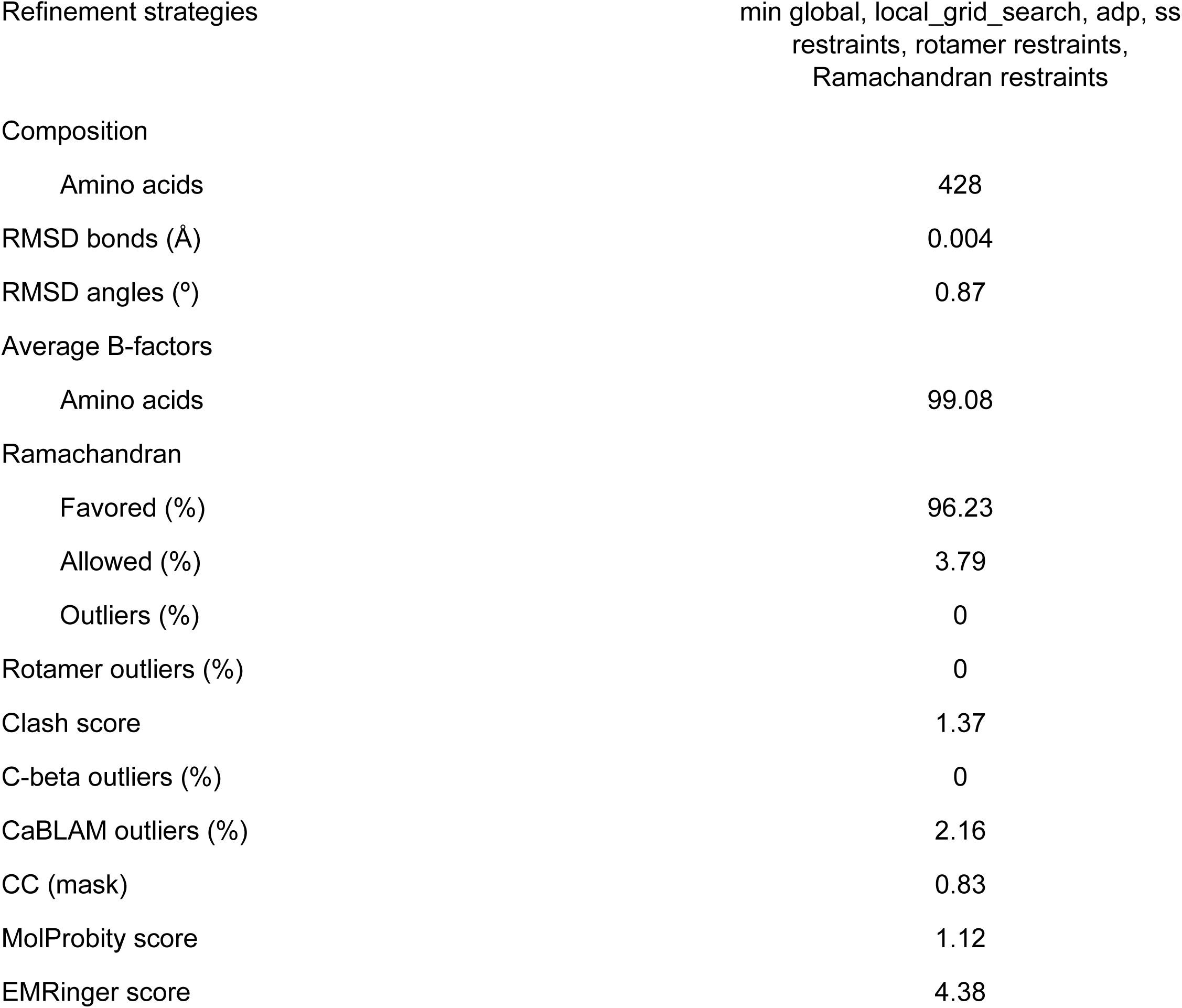
**PDB validation report**

---

**Supplemental Figure 1.**
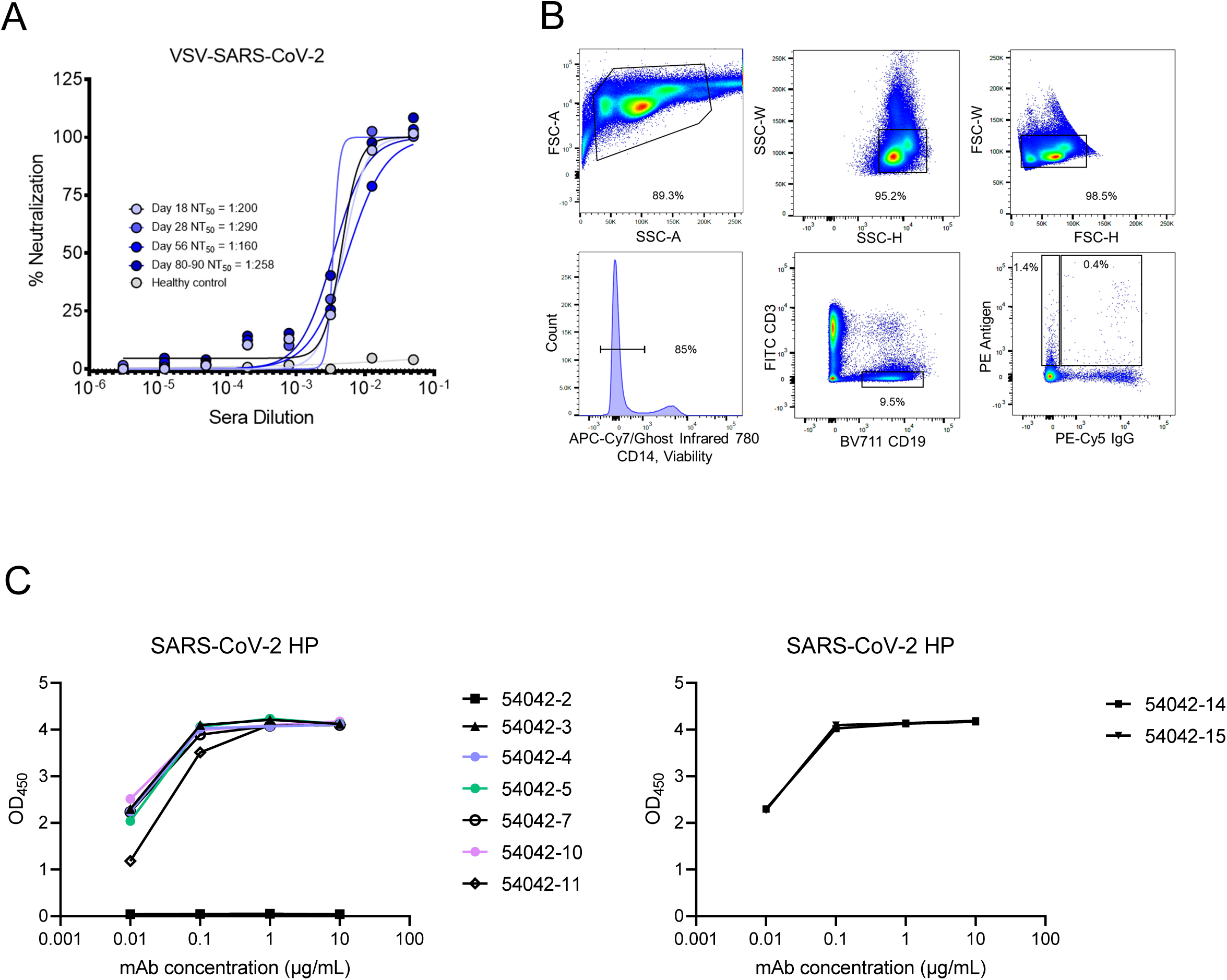
**(A)** VSV-SARS-CoV-2 capacity of serum is displayed from time points at day 18, day 28, day 56, and days 80-90. **(B)** Gating scheme for fluorescent-activated cell sorting of recovered COVID-19 individual. Cells were stained with Ghost Red 780, CD14-APC-Cy7, CD3-FITC, CD19-BV711, and IgG-PE-Cy5 along with a DNA-barcoded antigen screening library. To detect antigen-positive B cells, cells were washed and treated with a streptavidin-PE secondary stain. Gates as drawn are based on gates used during the sort, and percentages from the sort are listed. **(C)** ELISA binding data of candidate antibodies identified by LIBRA-seq against SARS-CoV-2 spike HP.

**Supplemental Figure 2.**
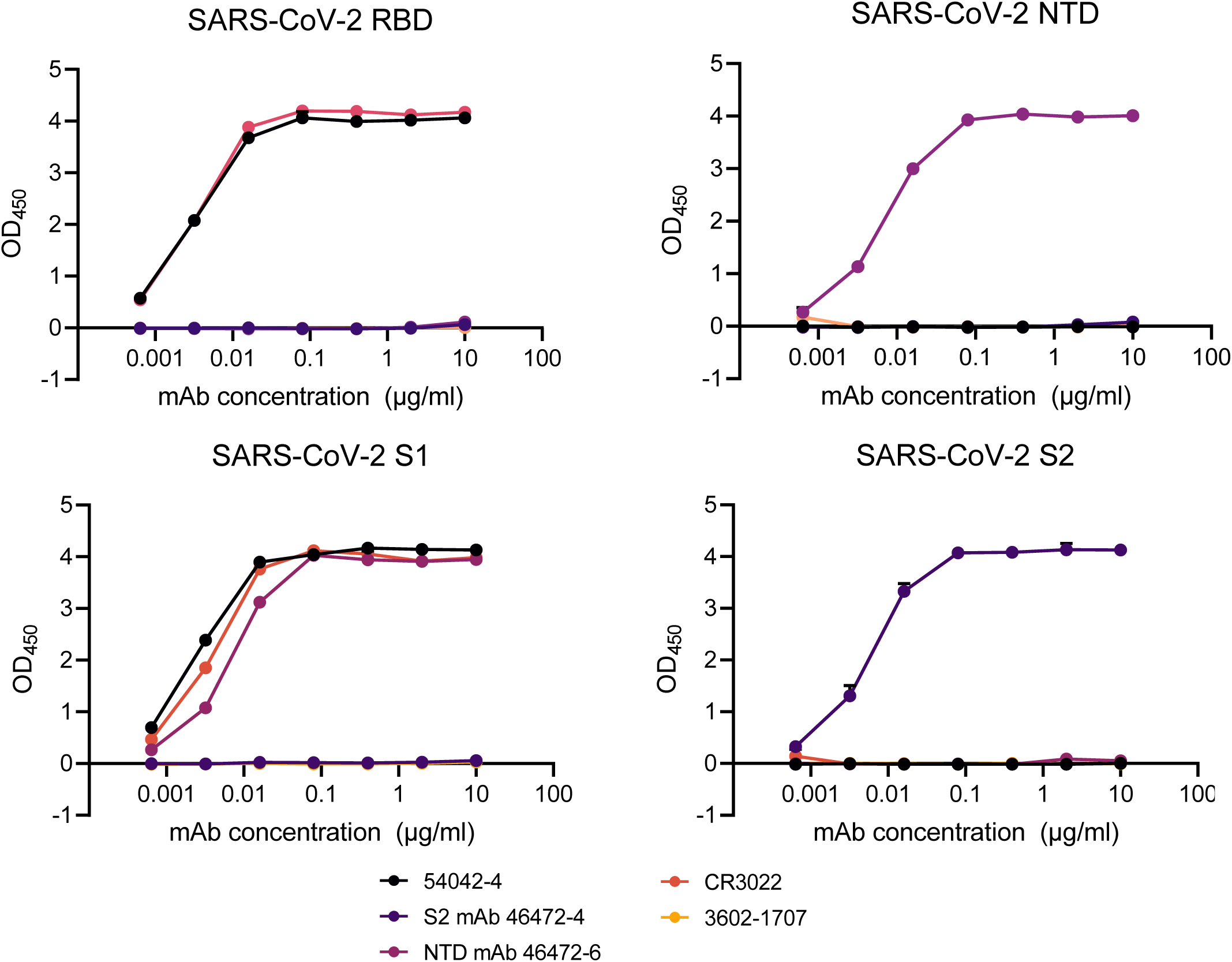
ELISA binding data against SARS-CoV-2 subdomains RBD, NTD, S1, and S2 are shown. CR3022 was used as a positive control RBD-directed antibody(Yuan et al., 2020a) whereas 46472-4 and 46472-6 antibodies were used as positive controls for the S2 and NTD, respectively (Shiakolas et al., 2020). The HA-specific 3602-1707 antibody was used as a negative control.

**Supplemental Figure 3.**
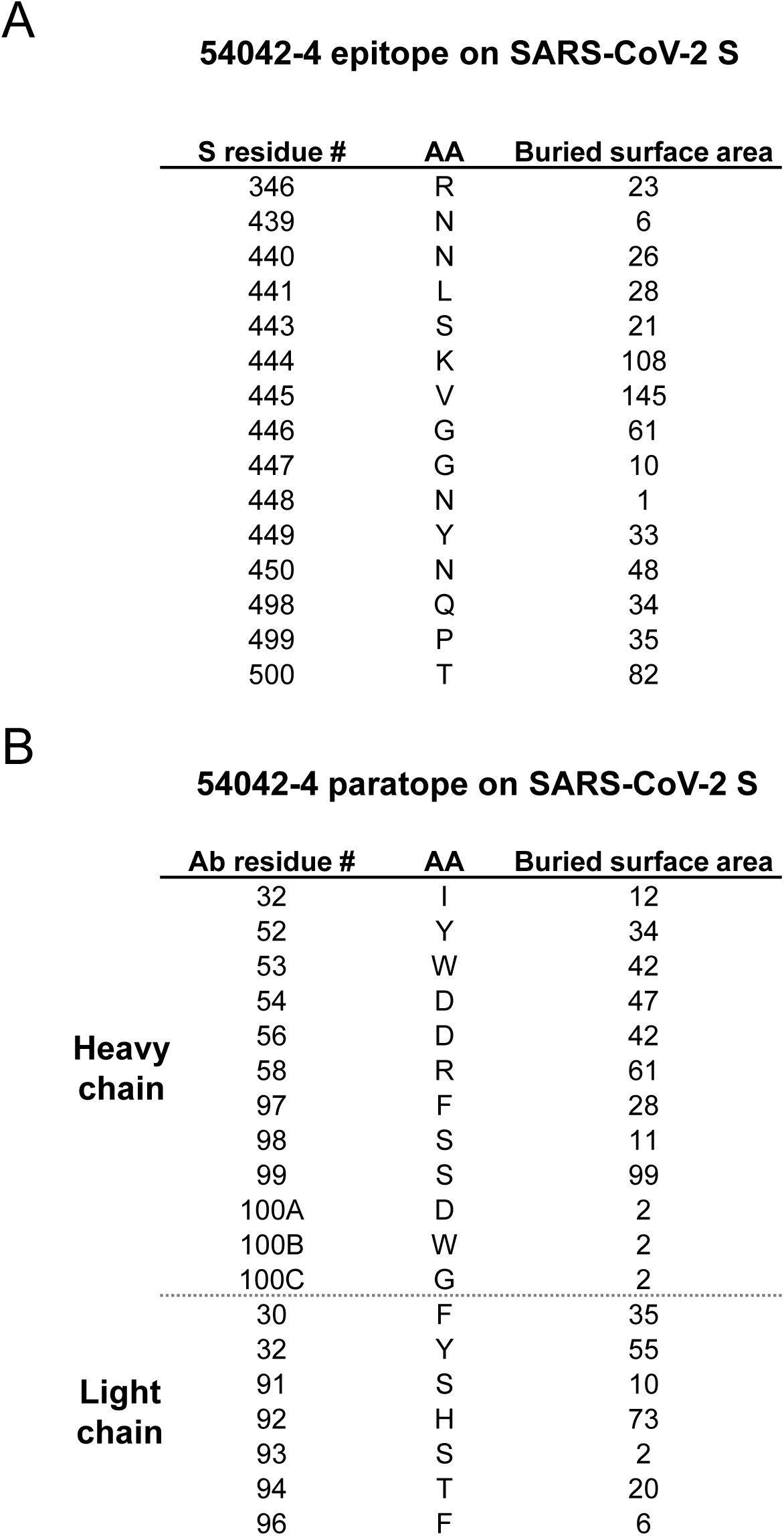
**(A)** SARS-CoV-2 spike residues comprising the epitope of 54042-4 are shown with their associated buried surface area (Å^2^). **(B)** 54042-4 residues comprising the antibody paratope against SARS-CoV-2 spike are shown with their associated buried surface area values (Å^2^).

**Supplemental Figure 4.**
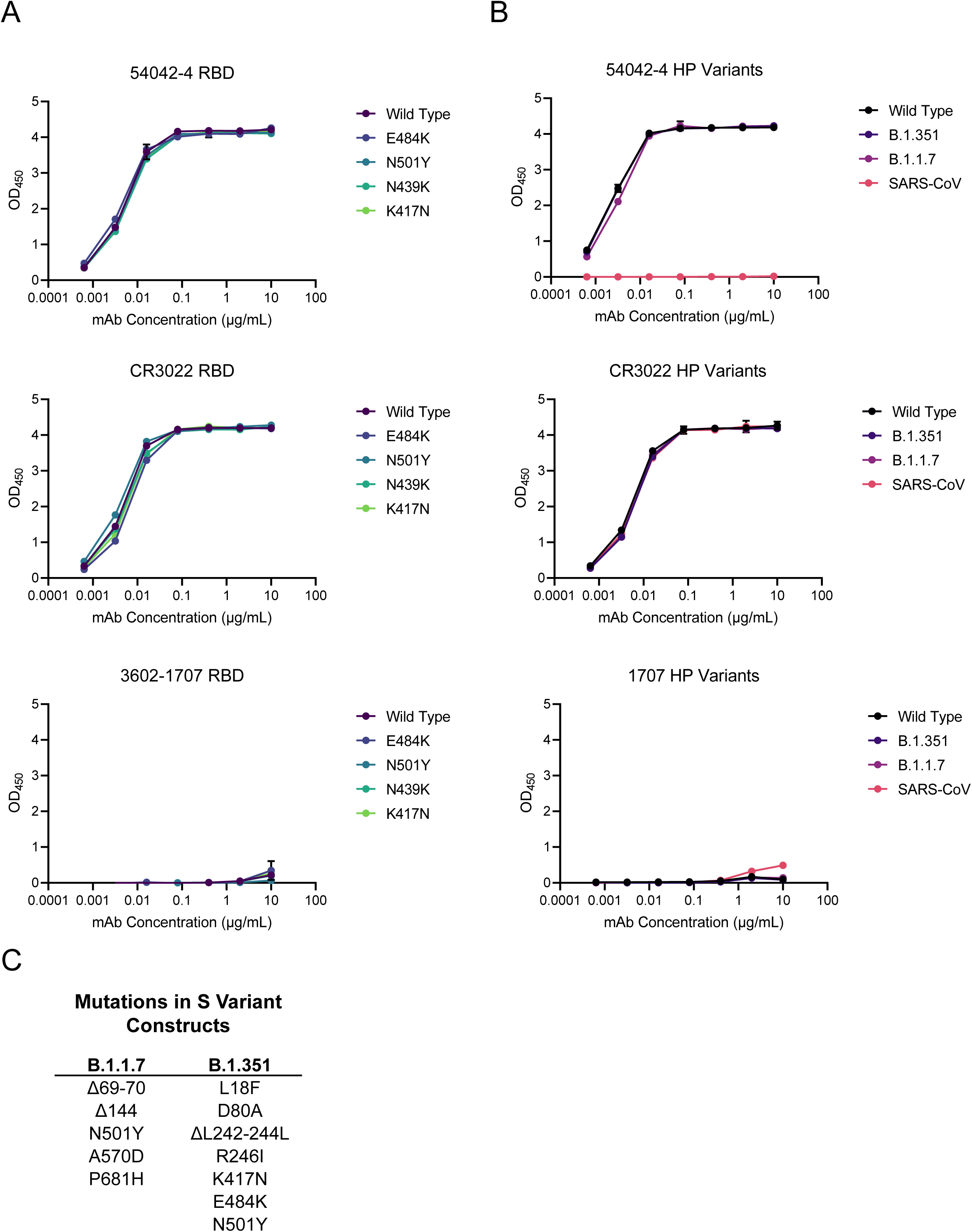
**(A)** ELISA binding data against SARS-CoV-2 Wuhan-1 RBD and RBDs with substitutions E484K, N501Y, N439K, or K417N. CR3022 was used as a positive control and 3602-1707, an HA-specific antibody, was used as a negative control. **(B)** ELISA binding data against SARS-CoV-2 S HP, SARS-CoV S, and SARS-CoV-2 S HP constructs with substitutions in the S1 domain for the B.1.351, and B.1.1.7 variants of concern. CR3022 was used as a positive control and 3602-1707 was used as a negative control antibody. **(C)** The substitutions and deletions present in the B.1.1.7 and B.1.351 constructs used in the ELISAs depicted in **Supplemental Figure 4B**.

**Supplemental Figure 5:**
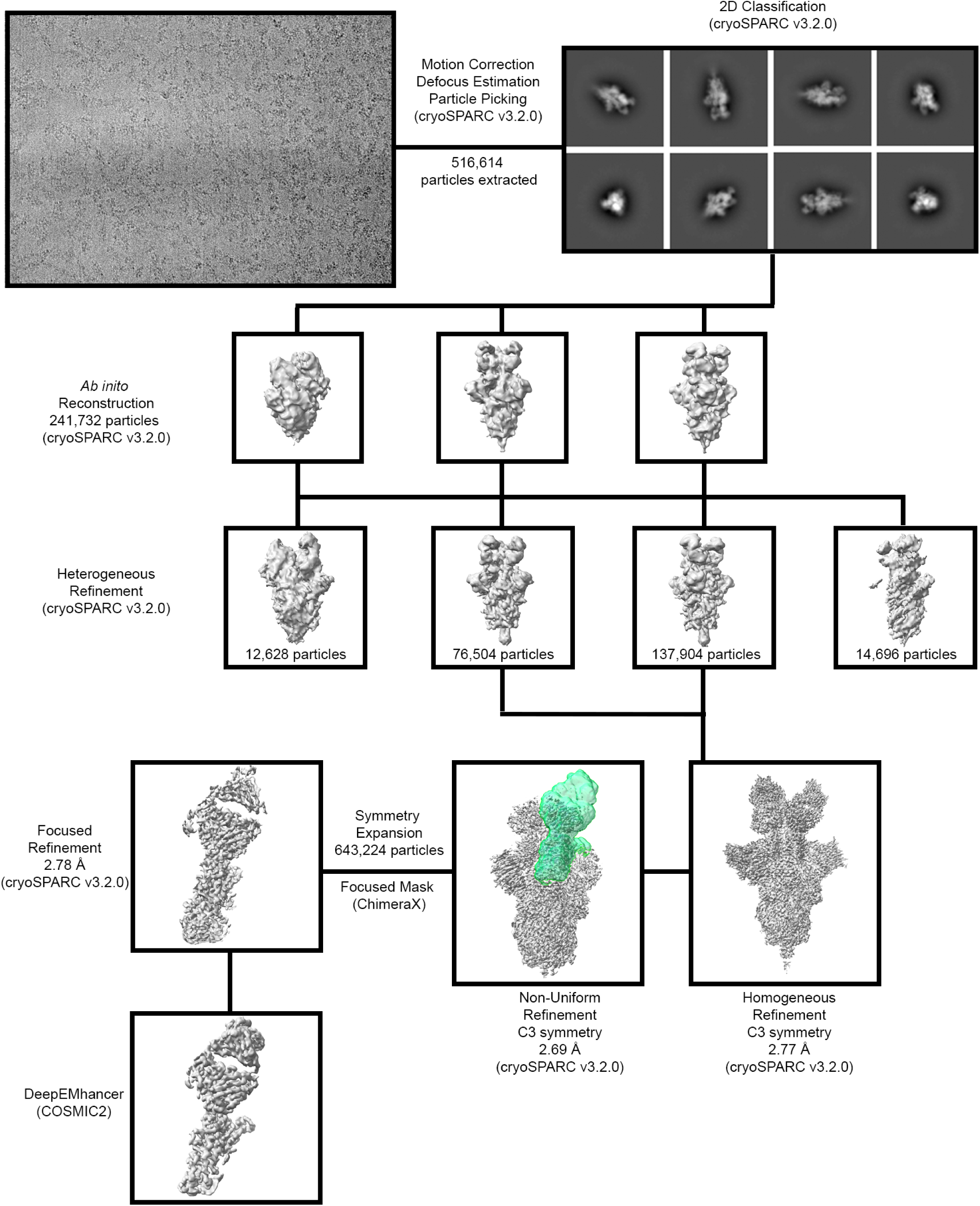
**Cryo-EM data processing workflow.** Flowchart outlining cryo-EM data processing of Fab 54042-4 Fab bound to SARS-CoV-2 S. Additional information can be found in the Methods section under “Cryogenic electron microscopy (cryo-EM)”.

**Supplemental Figure 6:**
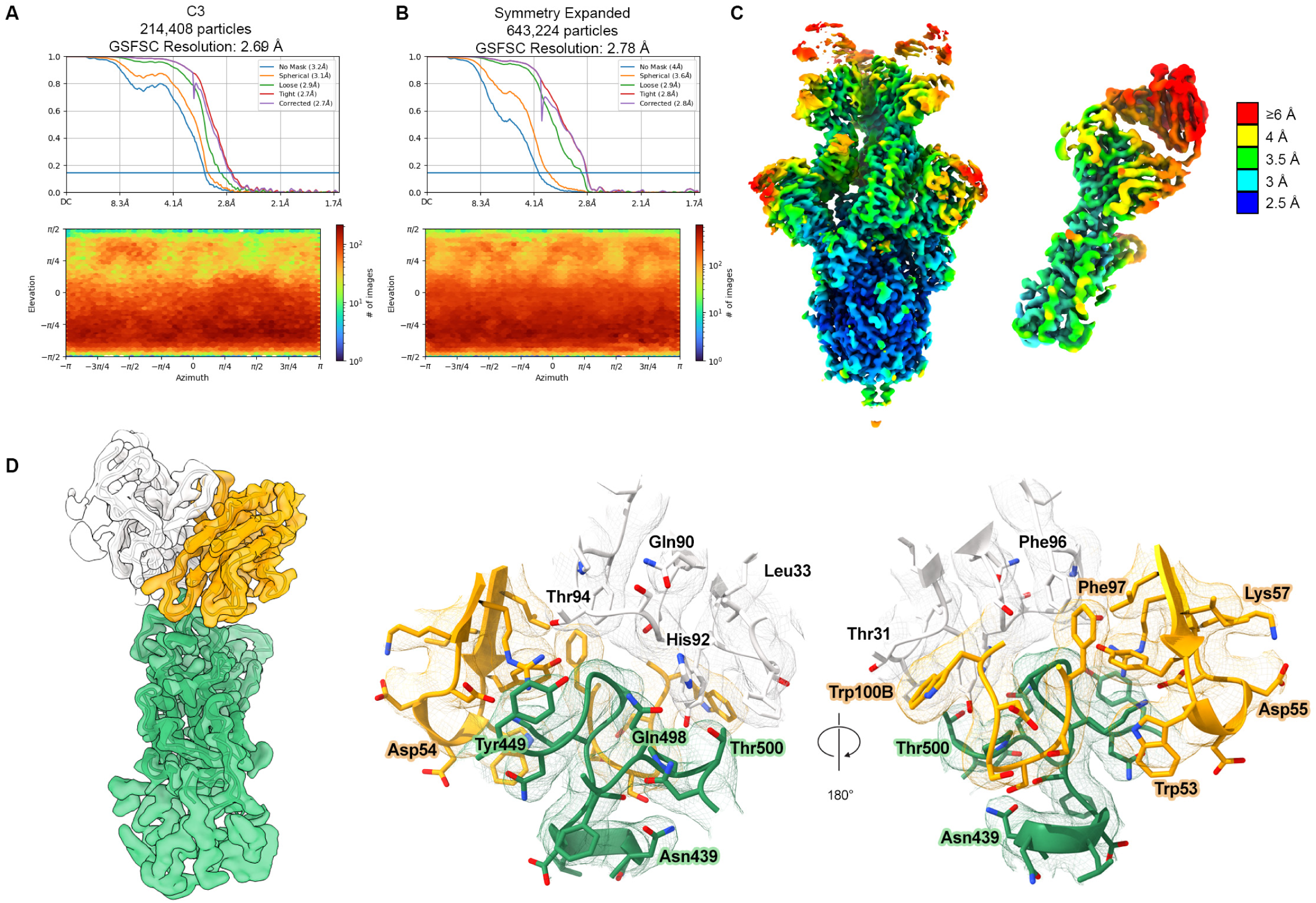
Cryo-EM structure validation. **(A)** FSC curve and distribution plot for the C3 S-ECD/54042-4 structure, generated in cryoSPARC v3.2.0. **(B)** FSC curve and viewing distribution plot for focused refinement of the S-RBD bound to 54042-4 Fab. **(C)** Local resolution shown by color of the C3 S-ECD/54042-4 (left) and focused S-RBD/54042-4 (right) reconstructions. **(D)** Map resulting from focused refinement of the RBD (green) (left), 54042-4 heavy chain (orange), and 54042-4 light chain (white). Detailed views of the binding interface and corresponding map (center, right). Oxygen atoms are colored red, nitrogen blue, and sulfur yellow.

